# miRXplain: explainable isomiR-aware microRNA target prediction using CLIP-L experiments and hybrid attention transformers

**DOI:** 10.64898/2026.02.11.703270

**Authors:** Ranjan Kumar Maji, Giulia Cantini, Hui Cheng, Annalisa Marsico, Marcel H Schulz

## Abstract

MicroRNAs (miRNAs) are ~22 nt long noncoding RNAs that repress genes by base-pairing with complementary sequences in target mRNAs. Variants known as isomiRs arise from alternative hairpin processing and often shift the canonical seed region, thereby altering their target repertoire. Yet, the mRNA features that enable differential miRNA and isomiR target selection remain poorly understood, largely due to limited high-throughput assays that capture the exact miRNA bound to each target. Existing deep learning methods for miRNA-target prediction have neither leveraged such datasets nor investigated isomiR-specific interactions. To fill this gap, we developed *miRXplain*, an isomiR-aware transformer that predicts miRNA–mRNA interactions using miRNA and target sequences derived from CLIP-L data, directly linking precise miRNA variants to their target sites. These data however revealed a 5′-end nucleotide bias in target sites, which we corrected to generate high-quality, miRNA–target pairs for training, preserving isomiR-specific signal. *miRXplain* outperformed all benchmarked models, surpassing TEC-miTarget, in auROC and auPRC with 15× fewer parameters. Attention maps highlighted distinct sequence determinants for canonical versus isomiR interactions, and *in silico* saturation mutagenesis highlighted the importance of seed and 3′ supplementary regions. *miRXplain* effectively prioritizes pathogenic single-nucleotide variants affecting mRNA–miRNA interactions and reveals isomiR targeting principles that deepen our understanding of miRNA biology.

## 1 Introduction

micro(mi)RNAs are ~22 nucleotide (*nts*) long non-coding RNAs that are key regulators gene expression and hence cellular function. These small RNAs bind to their target transcripts and thereby sculpt the transcriptome. The miRNA seed region, 2~7 positions from the 5′ end of the mature sequence, has been known to drive the selection of mRNA targets, their repression, and/or translation inhibition. However, studies have shown that the miRNA target interaction is mediated by many factors, such as hybridization of the miRNA target duplex, target site accessibility, and interactions with RNA-binding proteins, among others [1]. miRNAs can regulate targets through canonical seed-matched 3′UTR binding, as well as non-canonical interactions that involve imperfect seed pairing or binding at alternative transcript regions [2].

Under cellular stress conditions, primary/precursor miRNAs are alternatively processed during biogenesis, thereby generating sequence variations in the mature miRNA form [3] called isomiRs [4–6]. Though sequence modifications can occur at any position of the canonical miRNA, in this work we focus on isomiRs with addition or deletion of nucleotides at the 3′, 5′, or both ends of the canonical miRNA (templated isomiRs). IsomiR modification at the 5′ end shifts the miRNA seed region compared to its canonical form and results in the selection of different target sites. 5′ isomiRs are functional and are of evolutionary importance [7]. For example, canonical miR-411 and its 5′ isomiRs repress different target pools [8] contributing to different functions [9–11]. While numerous prediction tools have been developed to predict canonical miRNA–mRNA interactions, there is currently no *available* approach to systematically predict both canonical and isomiR–mRNA target site interactions.

Prediction of miRNA–mRNA interactions has traditionally relied on RNA–RNA sequence alignments (PicTar [12]), thermodynamic stability of miRNA–target hybrids (e.g. miRanda [13], RNAhybrid [14]), or the identification of conserved canonical seed matches in multi-species alignments, e.g. TargetScan [15]. Tools such as TargetScan were largely developed on sites with evolutionary conservation and experimental validation, whereas miRanda integrates sequence complementarity with hybridization free energy and was benchmarked against validated interaction datasets. Genome-wide miRNA target prediction can be refined by integrating cell type– or condition-specific miRNA and gene expression profiles [16–18]. Because these methods rely on strict assumptions, based on the most ideal required feature set for RNA interactions, they often miss real interactions or include biologically implausible ones, leading to false positives and false negatives.

Cross-linking immunoprecipitation sequencing (CLIP) [19] experiments typically use antibodies against the Argonaute (*Ago*) protein to isolate Ago-bound mRNA regions, as a proxy to identify miRNA targets. Since small RNAs such as miRNAs are known to primarily act as an *Ago* guide, these CLIP-derived binding sites have been hypothesized to correspond to miRNA target sites. In practice, this means that based on existing target prediction methods, a plausible miRNA is predicted that may have guided *Ago* to this position. CLIP-L methods such as CLASH (cross-linking, ligation, and sequencing of hybrids), [20], qCLASH (quick CLASH) [21, 22], CLEAR-CLIP [23], and related ligation-based CLIP methods, have extended conventional Ago-CLIP by introducing an RNA ligation step that covalently links the guide miRNA to its target fragment. This allows direct sequencing of small RNA-mRNA chimera, including miRNAs, providing nucleotide-resolution evidence of the interacting pairs and thus avoiding the need to guess the miRNA involved. Furthermore, CLIP-L not only captures the specific miRNA in the interaction, but in addition captures sequence variations of it, including isomiRs, improving miRNA–target site pair specificity. Published CLIP-L analysis pipelines such as *hyb* [24], *hybkit* [25], and SCRAP [26], have annotated only canonical miRNAs and other small RNA classes (such as tRNA, rRNA). They did not consider RNA sequence variations ligated to the target site in the chimeras.

Many machine learning (ML) methods, such as microCLIP [27, 28], have been trained on Ago-CLIP data, using experimentally derived binding regions as positive examples to improve miRNA–mRNA interaction prediction. Ligated CLIP experiments, in contrast to Ago-CLIP, are seldom utilized to develop miRNA target prediction models. While ML methods improved prediction by leveraging handcrafted combinations of miRNA and mRNA features [29, 30], they remain limited to predefined characteristics known to influence interactions. In contrast, deep learning (DL) approaches can extract complex sequence features directly from small RNA–target pairs, capturing interaction-specific patterns without relying on handcrafted inputs, and thereby outperforming classical methods.

Several DL models, including MirTarget v4.0 [31], miTAR [32, 33], TEC-miTarget [34], Mimosa [35], DMISO [36], and GraphTar [37] have mostly exploited Ago-CLIP datasets for training. However, as mentioned above, the need to predict the miRNA involved in an Ago-CLIP interaction means that all ML and DL methods trained on these data assume that this prediction is correct. In reality, the wrong miRNA may be included in the training set, for example when another small RNA is involved in the interaction [38]. This biases models towards predefined assumptions and limits the exploration of broader miRNA/isomiR biology, and as such the progress made by recent DL approaches is only incremental due to the limitations of Ago-CLIP data for training. The most recent transformer-CNN model, TEC-miTarget, which uses predicted miRNA and CLIP-derived target binding sites, outperforms other DL methods, but remains computationally expensive and was not tested for isomiRs. DMISO [36], is the only approach designed to predict canonical miRNA and isomiR targets; however, its implementation is unavailable. Thus, no tested isomiR-aware target prediction model currently exists.

To fill this gap, in this paper, we developed *miRXplain*, an efficient and explainable isomiR-aware DL model that leverages high-resolution CLIP-L data to learn RNA–RNA interaction specificities, correct sequence biases, and report miRNA–target interactions for both canonical miRNAs and diverse isomiR classes. We systematically process CLIP-L datasets to correct for ligation bias across experiments, enabling reliable identification of genuine canonical miRNA and isomiR sequences, along with the corresponding mRNA sequences at interaction sites, to train the model. Leveraging bias-corrected CLIP-L derived target sequences, *miRXplain* simultaneously learns miRNA/isomiRs sequence representations, target sequence representations, and RNA–RNA interaction specificities through a deep architecture combining Transformer encoders with convolutional neural network (CNN) layers. Trained on a large experimental dataset that includes isomiR interactions, this isomiR-aware model employs its multi-module architecture to deliver accurate miRNA target predictions, capturing interaction patterns for both canonical miRNAs and diverse isomiR classes. Crucially, unlike previous methods that either ignore or cannot accommodate isomiRs by design, *miRXplain* can be explicitly questioned on rules of isomiR biology using explainability approaches. We show that *miRXplain* consistently outperforms state-of-the-art architectures across multiple metrics for both canonical miRNA and isomiR–target interaction prediction. *miRXplain* is computationally efficient and thus opens up a new path to explore large-scale, miRNA and isomiR target interactions.

## 2 Methods

### 2.1 IsomiR-aware processing of CLIP-L experiments

IsomiRs, variants of the canonical miRNAs, have nucleotide modifications compared to the mature canonical miRNA form (**Fig.** 1A). Their interactions with the mRNA target site can be retrieved from CLIP-L experiments. These experiments generate chimeric reads that comprise of miRNAs or isomiRs (small RNA region), tethered with their mRNA target sites (**Fig.** 1B). To annotate the miRNAs and their isomiRs in the chimeric reads from CLIP-L experiments (**Table ??**), we adapted the SCRAP pipeline, originally developed to annotate CLIP-L chimeras with known small RNAs and transcripts [26] (**Supplementary Fig.** 1). The adapter sequences and barcodes were trimmed from the chimeric reads using Cutadapt v4.4 [39] and trimmomatic [40]. We aligned the chimeric reads to canonical miRNAs and their corresponding hairpin sequences from miRBase [41]. If the small RNA region of the chimeric read aligned completely with the canonical miRNA sequence, it was annotated as a canonical miRNA. The small RNA region was annotated as a variant of a specific canonical miRNA, or isomiR with an addition event, when the read alignment spanned the canonical miRNA and extended to align the associated hairpin sequence. If the small RNA region of the chimera aligned to the hairpin with an addition of *x* nts on either end (say the 5′ end) relative to the canonical miRNA, then ‘|5′ + *x*’ was added to the miRNA annotation, an example of a 5′ isomiR *addition* event. If instead the small RNA region aligned with a few nts less (say *y* nts) than the canonical miRNA, then the small RNA was annotated as ‘|5′ *x*’ isomiR, representing a *deletion* event. The remaining of the chimeric reads were next aligned to the 3′ UTR of the mRNA to get the target site on the transcript for the miRNA or isomiR. miRNAs and their target site sequence pairs were then used to train the classification model. GENCODE hg38 v44 was used for transcript annotations. A minimum target length of 20 *nts* was set to filter out target sites that potentially contribute to unstable hybrids (**Supplementary Fig.** 2).

**Fig. 1:**
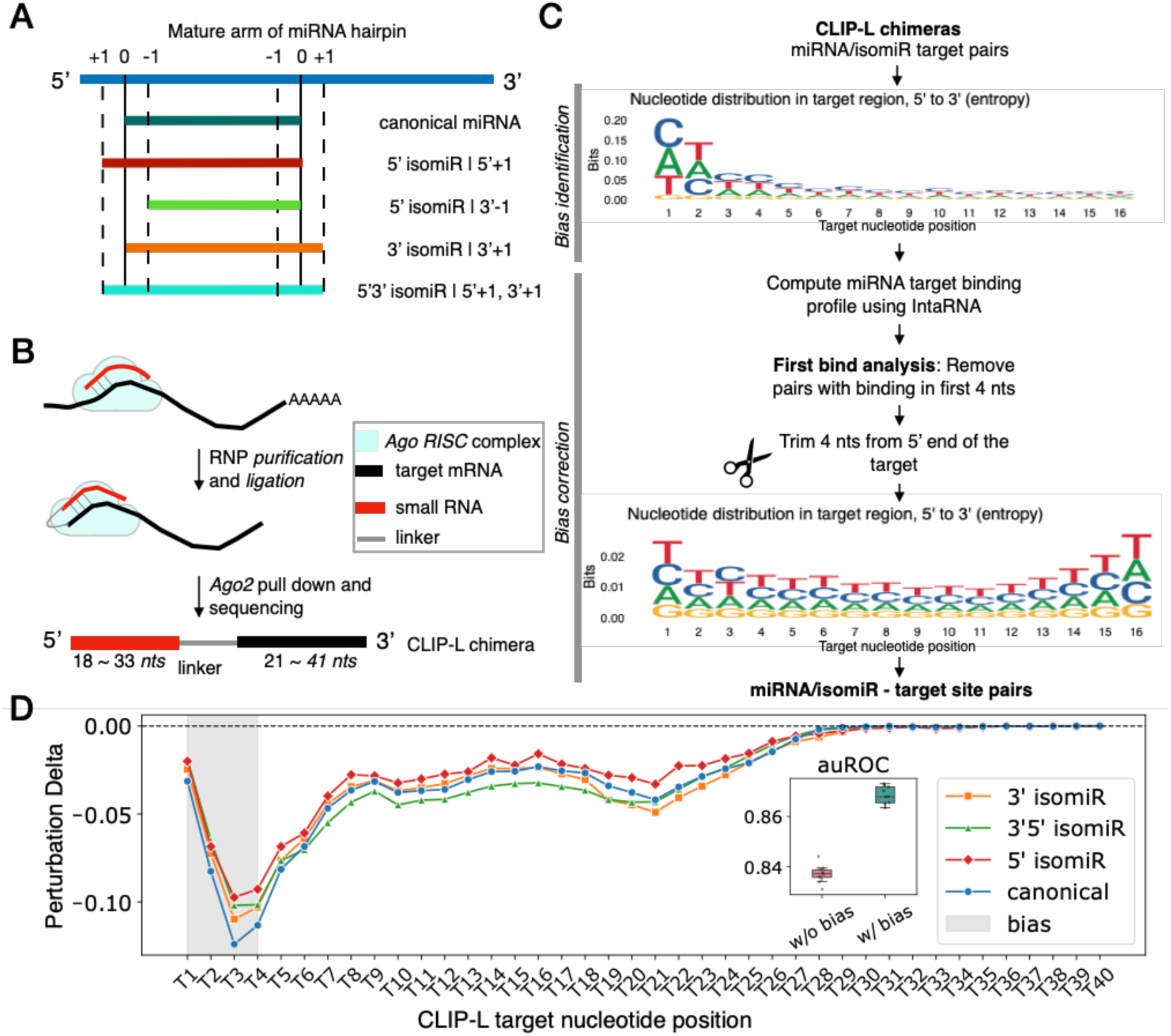
CLIP-L chimeras without addressing biases do not reveal miRNA and target binding variations. **(A)** Definition of isomiRs of its canonical miRNA forms as considered in the study. A few examples of the various isomiRs with addition and deletion events (or tailing and trimming respectively), and their representative annotations are shown. **(B)** Schematics of the CLIP-L protocols, with the ligation step that tethers the small RNA guide to the target mRNA sequence. The Ago2 pull-down followed by sequencing generates the CLIP-L chimeras, with the small RNA and the target site sequence. Processing the chimeras can reveal the miRNA and its variant that interacts at the target site. **(C)** Workflow to identify and filter out biased CLIP-L interactions. The example biased sequence logo in the workflow corresponds to CLIP-L target sites from Kozar et al. **(D)** CNN*-baseline* model interpretation on the biased dataset highlights that the model learn only the biases, and not the miRNA target rules for canonical and isomiR types. The inlay shows the difference in model performance using miRNA and target pairs before (w/ bias) and after bias correction (w/o bias).

### 2.2 Bias identification and correction in the CLIP-L datasets

Minimum Free Energy (MFE) for miRNA and their targets from the CLIP-L chimeras were computed using RNAcofold [42] to assess whether they captured unstable miRNA-target duplexes (**Supplementary Fig.** 3). We examined the miRNA target sites identified by the CLIP-L experiments within transcript 3′ UTRs, extracted from the chimeras as described in the previous section, to assess potential sequence bias introduced by the protocols. For each protocol, we generated sequence logos of the target sequences from the CLIP-L–derived chimeras (**Supplementary Fig.** 4). This analysis revealed nucleotide bias at the 5′ ends of the targets (**Fig.** 1C).

To further investigate whether these biases were influencing the model performance and hindering interpretation, we trained a simple convolutional neural network to predict miRNA-mRNA interactions, hereafter referred to as CNN*-baseline* (**Supplementary Fig.** 5) on two sets of the data, one retaining the bias and one with the bias removed (described in next section). Thereafter, we performed *perturbation analysis* on the CNN*-baseline* to confirm that the biased sequence positions on the target influenced the predictions of the model (**Fig.** 1D).

**Table 1:**
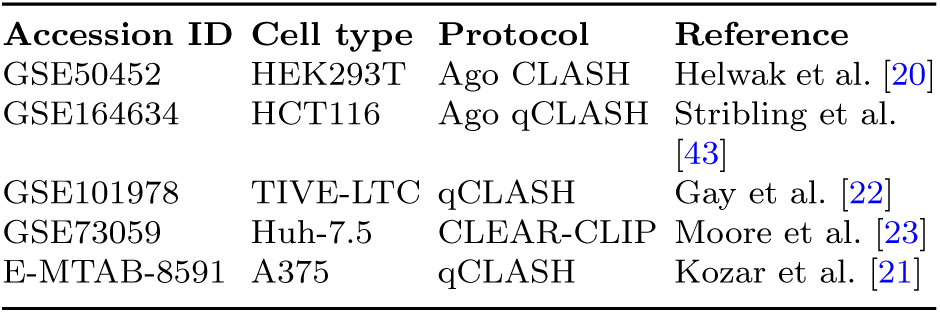
Human CLIP-L experiments used for training *miRXplain*.

### 2.3 Curation of positive miRNA target sites

To account for the nucleotide biases found in the CLIP-L experiments, we first predicted binding specificity with IntaRNA [44] using parameters: –seedBP 4 (number of base pairs within the miRNA seed) as the seed constraint (**Fig.** 1C). Interactions with miRNA target binding within the first 4 *nt* positions were removed, while retaining the first binding position of the miRNA over the target within 12 nts from the 5′ side, based on previous observations from RBNS experiments [45]. Furthermore, to compensate for the loss of target context caused by trimming, and to evaluate whether additional flanking information improves model performance, we extended the target sequences in increments of 5 nucleotides at the 3′ end. The optimal length of the target sequence was determined to be 5 additional *nts* by training the CNN*-baseline* model on trimmed datasets (**Supplementary Fig.** 6). This constituted bias-corrected positive training examples for our model training and evaluation.

### 2.4 Generation of negative miRNA target sites

To build a balanced training dataset for the classification task, we generated a negative sample for each positive instance. We sampled the potential negative sites from neighboring 3′ UTR regions for each positive miRNA–target pair, ensuring that none contained binding sites of any other miRNA across all experiments (**Supplementary Fig.** 7). To reduce the risk that the model exploits potential GC-content biases in positive interactions, candidate negative sites were selected to match the GC-content of the corresponding positive interaction within a tolerance of ±0.01. There is no ground truth data to ascertain that candidate negative sites are not potential miRNA interaction sites. MFE being an important factor that drives miRNA binding at the target site [33], we used MFE as a filter to select potential unstable interactions among candidate negative sites. Thus, from the selected candidates, sites with the highest MFE were chosen as the negative sample. This stringent sampling strategy allowed us to define plausible negative sites which, when compared to positive sites during training, enabled the model to focus on the salient features of miRNA-target binding. The final balanced training dataset contained a total of 102,872 miRNA–mRNA interaction pairs, with 53,386 (52%) positive samples and 49,486 (48%) negative samples.

### 2.5 miRXplain architecture

*miRXplain* is a sequence-based Transformer network that employs a hybrid attention mechanism to predict binding probability for miRNA/isomiR and target sequence pairs. The overall architecture (**Fig.** 2A), in addition to the sequence *embedding layers*, comprises of hybrid self- and cross-attention between miRNA and its target sequences (**Fig.** 2B), feature fusion module, the convolutional module for refining interaction features (**Fig.** 2C) and finally, the classification module (**Fig.** 2D).

**Fig. 2:**
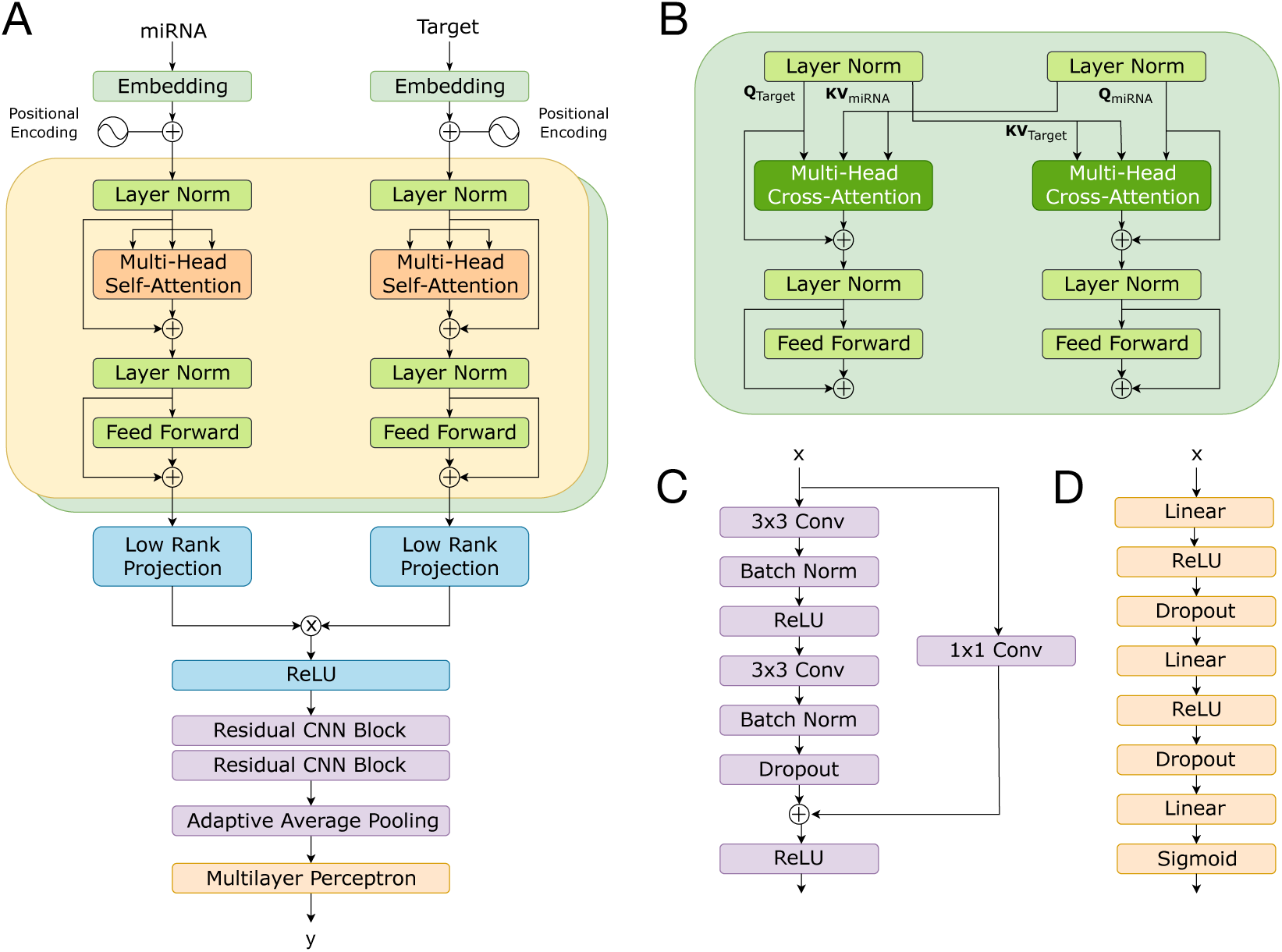
Overview of the *miRXplain* architecture. **(A)** Overall model architecture displaying separate input embeddings for miRNA and target with positional encoding, Transformer encoder layers with hybrid attention and low-rank fusion module followed by a convolutional block and MLP. **(B)** The second layer of the hybrid attention mechanism illustrating cross-attention between miRNA and target sequences. **(C)** Residual CNN block for feature refinement. **(D)** Final classification multilayer perceptron (MLP).

### (i) Encoding module

#### Padding and embedding

Since miRNAs, isomiRs, and their corresponding target sites (as captured by the chimeric reads) can vary in length, we uniformed the sequence lengths within each branch by padding as needed before inputting them into the Transformer encoder. Taking into account the distribution of sequence lengths occurring in our dataset (**Supplementary Fig.** 8), the sequence nucleotides of the miRNA/isomiR and the target site pairs were padded to a maximum length of 33 and 41 *nt* s respectively, using the unknown nucleotide N. After padding, sequences were converted to an integer representation (*N-0, A-1, U-2, C-3, G-4)* and separately fed to an *embedding layer* mapping discrete inputs to dense vectors of size *d_model_*, allowing the model to learn semantic similarities between nucleotides.

#### Positional encoding

To encode positional information in the input sequences, we followed the standard positional encoding implementation introduced in the original Transformer [46]:

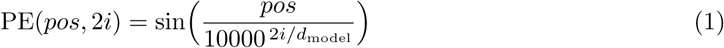

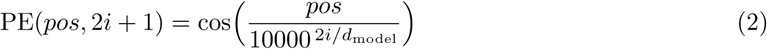

where *pos* is the position index and *i* is the dimension index. Additionally, we tested an alternative mechanism that included a learnable scalar weight per position; however, this did not yield performance improvements over the standard configuration. The positional encodings and embedding vectors are then summed and passed as input to the next module.

### (ii) Feature extraction module

To capture *intra*- and *inter* -sequence dependencies between miRNA and mRNA target site, a hybrid attention mechanism was employed, with self-attention followed by cross-attention layers (**Fig.** 2). Self-attention is calculated independently on the miRNA and the target branch, following the formula:

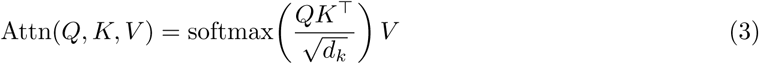

where 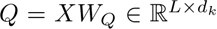 is the query, 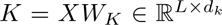 the key and 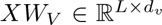 is the value vector, 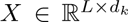 is the input miRNA or the target sequence and 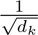 is a scaling factor used to stabilize the gradients during training.

In *self-attention Q*, *K* and *V* are all derived from the same sequence i.e. the miRNA or the target. In contrast, *cross-attention* is implemented to enforce the model to learn interaction information to flow from the miRNA to the target and vice versa, thus allowing each miRNA position distribute attention over target positions (*Q*_miRNA_*, K*_target_*, V*_target_)

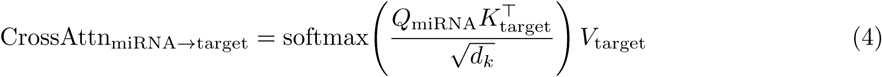

thereby making each target position attend over the miRNA sequence (*Q*_target_*, K*_miRNA_*, V*_miRNA_)

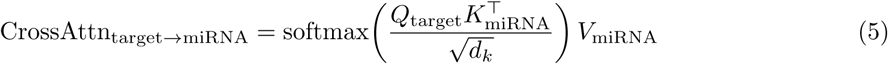

Following the attention layers, a position-wise Feed Forward Network (FFN) with GELU (Gaussian Error Linear Unit) activation transforms each token independently to build non-linear features.

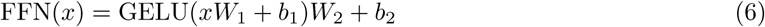

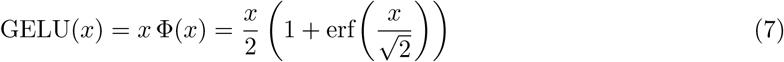

where Φ(*x*) is the standard normal cumulative distribution function (CDF), and *erf* (*u*) = erf(*u*) = 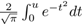 is the error function.

To stabilize training, the model additionally incorporates residual connections and layer normalization before the attention and FFN blocks, as:

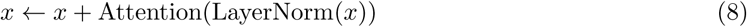

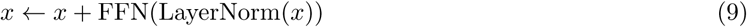

Furthermore, dropout is applied within both attention and feed-forward blocks to regularize training and improve model generalization.

### (iii) Feature fusion module

To efficiently integrate features from miRNA and target, *miRXplain* employs the Low-Rank Bilinear Fusion (LBF) approach, originally developed for multimodal representation learning [47]. LBF models pairwise interactions between two different vectors. Given two input vectors *x* ℝ*^d^* and *y* ℝ*^d^*, the standard bilinear fusion can be expressed as:

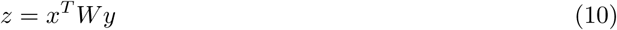

where *W* ℝ*^d×d^* is a weight matrix, modeling the second-order interactions between the two vectors. To optimize this computation, potentially expensive in case of high input dimensions, LBF introduces an improvement by approximating *W* with two low-rank projection matrices *U, V* ℝ*^d^*^×*r*^, where *r<<d*. The interaction is then computed as:

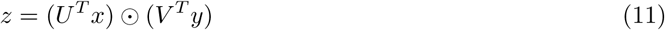

where Θ denotes element-wise multiplication operation. This low-rank factorization reduces the number of parameters from (*O*(*d*^2^) to (*O*2*dr*), significantly improving efficiency while maintaining sufficient expressiveness after tuning the matrix rank *r*.

As in the previous module, layer normalization is applied to both projected vectors before the fusion. A ReLU activation was applied afterwards to introduce non-linearity and therefore boost model capacity. The final computation is then:

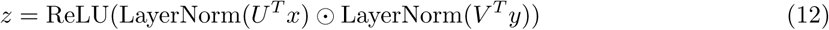

### (iv) CNN refinement module

For each pair of miRNA and target mRNA, the LBF module produces an output tensor of shape (*r, L_m_, L_t_*), where *r* is the low-rank dimension, *L_m_* and *L_t_* are the lengths of the miRNA and target sequences, respectively. This tensor can be interpreted as a two-dimensional “interaction map” where each channel represents a distinct interaction pattern between the miRNA and target mRNA sequences. To further model local patterns and spatial dependencies within this interaction map, *miRXplain* employs a CNN-based refinement layer.

Specifically, two residual CNN blocks are stacked, each consisting of a two-layer convolution architecture with 3×3 convolutions and padding of 1 to preserve spatial resolution, followed by batch normalization and a ReLU activation. Dropout is applied after the second convolutional layer to randomly discard a portion of the channel activations, improving the generalization and preventing overfitting while at each block we also include skip connections to facilitate gradient flow and mitigate the vanishing gradient problem.

Lastly, to address the variability in input sequence lengths, the model employs adaptive average pooling to compress the output interaction maps into fixed-size maps by dynamically adjusting the pooling window according to the input dimensions. The resulting maps are then fed to the classification module for output prediction.

### (v) Classification module

In this final module, the pooled features are flattened and passed through a three-layer perceptron equipped with ReLU activation and dropout for regularization before classification. The output layer employs a softmax activation to produce the final class probabilities.

### 2.6 Training setup

For training we utilize Adam [48] as optimizer with an initial learning rate of 0.0001 and batch size set to 32. These values were determined through hyperparameter tuning (see next section, *Tuning hyperparameters*). *miRXplain* was implemented in PyTorch Lightning v2.5.5 [49] and trained using 10-fold cross-validation, with 81% of the data for training, 9% for validation, and 10% for testing in each run. For every split, the proportion of positive to negative samples was preserved to match the distribution in the total dataset.

A learning rate scheduler is employed to reduce the learning rate by a factor of 0.2 if the validation loss does not improve for 3 consecutive epochs. In addition, L2 weight decay with a coefficient of 1×10^↘4^ is applied to all trainable parameters. To avoid overfitting and optimize computation time, early stopping was implemented by monitoring the validation loss, and training terminated if the loss did not improve for 5 consecutive epochs.

### 2.7 Tuning hyperparameters

Hyperparameter tuning was performed using grid search with the *Optuna* optimization framework v5 [50]. The search space explored was generated through combinations of the values reported in **Supplementary** **Table** 1. The optimal model hyperparameters were determined by evaluating each candidate configuration on 3 out of 10 cross-validation folds and selecting the configuration with the lowest mean test loss (**Supplementary Table** 2).

### 2.8 Benchmarking setup

Published DL approaches for miRNA target prediction have used CLIP data with fixed length CLIP-target sequences and considered predicted canonical miRNA sequence with limited length variability compared to isomiRs. *miRXplain* was benchmarked against DL models of various architecture types for miRNA-target prediction, specifically (i) two hybrid convolutional recurrent networks: *miTAR* and *DMISO*, (ii) a graph neural network based on a *word2vec-like* encoding: *GraphTar*, (iii) transformer-based architectures: *Mimosa* and *TEC-miTarget*, and finally a convolutional network, the CNN*-baseline* (**Fig.** 1D, **Supplementary Fig.** 5). These DL models have used fixed target input sequence lengths from Ago-CLIP data (such as 40 nt targets as processed in miRAW), however miRNA target pairs from CLIP-L data consist of variable miRNA/isomiR and target sequence lengths. To overcome the implementation heterogeneity, with their distinct strategies for feature extraction, training, data splitting, and model evaluation, we re-implemented their architectures to ensure a fair benchmark of recent DL methods. We created a unified framework for training and evaluation with consistent cross-validation splits, optimization strategies across different architectures using *PyTorch Lightning*. This process was straightforward for models already implemented in PyTorch, such as *GraphTar*, *TEC-miTarget*, and *Mimosa*, but required full porting across DL frameworks for *miTAR* and a complete re-implementation of *DMISO*, as no public code defining its architecture was available. All benchmarked models were trained using the hyperparameter configurations as in their original code or manuscript, and were trained and evaluated on the same data split, as defined in the section above, *Training setup*. All experiments were performed with NVIDIA A100 with 40GB memory.

### 2.9 Performance evaluation metrics

We evaluated all models using two standard classification metrics, the Area Under the Receiver Operating Characteristic Curve (auROC) and the Area Under the Precision-Recall Curve (auPRC). Given the true positive rate (TPR), and the false positive rate (FPR) as:

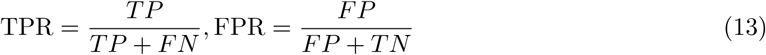

the ROC curve is obtained by plotting TPR against FPR across thresholds. The area under this curve is

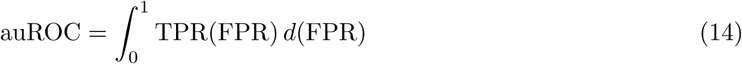

which represents the probability that a randomly chosen positive instance is ranked above a randomly chosen negative instance. This represents the ability of the classifier in distinguishing positive from negative instances. The precision and recall are given as:

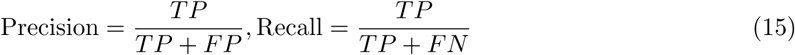

The precision–recall curve is obtained by plotting precision as a function of recall across decision thresholds. The area under this curve is:

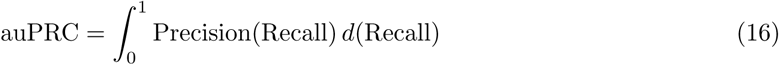

which quantifies average precision across all recall levels, emphasizing performance on the positive class under imbalance.

In addition to auROC and auPRC, we further assessed the performance of all models using accuracy, precision, recall and F1 score. Definitions of these metrics are given in **Supplementary Table** 3.

### 2.10 Background corrected self-attention maps

To probe which parts of the input miRNA sequence the model attended to during training, we visualized the self-attention weights of the miRNA encoder as square heatmaps. Each element of the heatmap is given by:

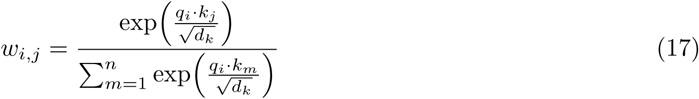

where, *w_i,j_* is the attention weight between query position *i* and key position *j*; *q_i_*: query vector at position *i* (row index); *k_j_*: key vector at position *j* (column index); *d_k_*: dimensionality of the key vectors (used for scaling), expanding on **Eq.** 3, with *Q* and *K* being computed over the miRNA sequence.

We implemented background correction to eliminate spurious correlations that may be present in the attention maps, following [51]. To estimate the background signal, we trained a control model with approximately 50% classification accuracy on data for which the original class labels had been randomly shuffled. This model was then used to extract *random* self-attention maps, containing the background noise and devoid of any real biological signal. For each sequence, these random maps were subtracted from the raw attention maps to obtain *contrast maps* that were subsequently used for visualization. The contrast attention scores are thus defined as:

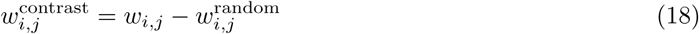

### 2.11 *In silico* saturation mutagenesis analysis

To investigate which regions of miRNA and target sequences contained the salient features used by the prediction model, we performed *in silico* saturation mutagenesis (ISM) analysis. by introducing nucleotide substitutions, this systematically tests all possible mutations at specific positions in the RNA sequence, to reveal how each change affects the model’s probability.

From 10 cross-validation models, we randomly selected a *miRXplain* model and its corresponding true positive test sequences with high prediction probability *p* 0.9 (from a total of 2,983 sequences). Each base (or *ref*) of the reference miRNA and target sequence (e.g. ‘A’) was mutated with other possible bases i.e., ‘C’, ‘U’, and ‘G’ (or *alt*). This yielded alternative pairs (in total 430,540) with a mutation present either in the miRNA or the target gene and were evaluated separately. The resulting mutant dataset was provided to the model to predict binding probabilities, and the single nucleotide mutation effects were evaluated by computing changes in predictions as

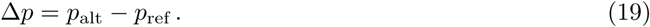

A negative or positive Δ*p* would imply a *loss-* or *gain* of binding probability through mutations, respectively.

### 2.12 Perturbation analysis

An additional method to quantify the importance of every nucleotide in the miRNA and target sequence is the *perturbation analysis*. This method evaluates single-nucleotide importance by masking bases with ‘N’, instead of mutating to other possible bases in each nucleotide position of the sequence (ISM), thereby effectively masking the original nucleotide identity from the model.

### 2.13 Analysis of canonical miRNA and isomiR targets

To characterize the potentially different target repertoire of isomiR and canonical miRNA target genes with our *miRXplain* model, we followed the workflow in **Fig.** 6A. We downloaded 3′ UTR sequences from UTRdb 2.0 [52]. The longest UTR for each gene was selected and binned into non-overlapping windows of length 30 *nt*. Remaining bins of less than 25 nts were removed. *miRXplain* was then used to predict target binding probabilities for each bin for the miRNA/isomiR of interest. A bin in the gene 3′ UTR was called a target bin with the classification threshold of *>*0.9, to maximise precision based on the PR curves (**Supplementary Fig.** 16). Significance of miRNA/isomiR binding to the 3′ UTR of a gene was obtained through binomial testing. The hypothesized probability of success in binding to the 3′ UTR bin for the miRNA of interest over the entire UTRome is given by:

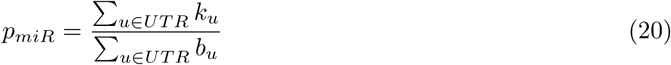

where *k_u_* is the number of bins where the miRNA *miR* is predicted to bind to UTR *u*; *b_u_* denotes the total number of bins in UTR *u* and *UTR* denotes the set of all 3′ UTRs considered for the analysis. Then the probability of observing a certain number of miRNA binding sites in a UTR can be computed with the binomial distribution:

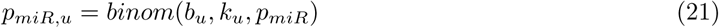

plugging the estimate *p_miR_* over all UTRs (**Eq.** 20). A one-sided binomial test (greater) is used to calculate the *p*-value for obtaining at least the observed number of miR hits in the given UTR. *p*-values are corrected for multiple testing (Benjamini-Hochberg) and individual genes are considered to be targeted in its UTR by a miR if the adjusted *p*-value 0.05. Thereafter, gene set enrichment analysis was performed on the top 500 target genes of the miRNA*/*isomiR of interest, ranked by their *p*-values, using R package *clusterprofiler* [53].

To evaluate if *miRXplain* predicted targets could recall experimentally validated miRNA targets, we curated reported interactions from miRTarBase 2025 [54] and DIANA TarBase-v9.0 [55]. We performed an unpaired Wilcoxon rank sum test of the miRNA predicted target genes ranked by their *p*-values compared to non-target genes.

### 2.14 SNV analysis

To evaluate SNV effects on miRNA binding, we obtained GWAS summary statistics from the UK Biobank [56] and filtered for significant SNVs (1% FDR). The SNVs were then compared to the miRNA genomic coordinates (hg19) from miRBase v21 [41]. To evaluate the effects of SNV mutations on miRNA binding, the binding probability was computed over the 3′ UTR of the target genes of experimentally validated miRNA targets (**Eq.** 21). To evaluate the significance in the difference of the binding probabilities, we performed a paired Wilcoxon rank sum test over the ranked *p*-values of all validated target genes for the mutated and the canonical (no mutation) miRNA sequence.

To analyze the effect of mutations for any nucleotide and miRNA position compared to the allele at the SNV position, the targets of the miRNAs were obtained from the CLIP-L experiments. The nucleotide at the SNV position on the miRNA (e.g., an A) was substituted with each other possible nucleotide (C, G, and U) and similarly, mutations were also introduced at any other position within the miRNA. We then used *miRXplain* to predict the binding probability for both the original SNV-containing sequence and all sequences carrying these artificially introduced mutations. Δ*p* (**Eq.** 19) was computed for each alternative nucleotide of the miRNAs that overlapped SNVs. The SNP IDs were obtained from NCBI dbSNP [57].

## 3 Results

### 3.1 CLIP-L captures genuine transcriptome-wide interactions of miRNAs and isomiRs with their targets

CLIP-L experiments that use an *Ago2* antibody for the Ago complex pull-down, capture specific small RNAs that guide *Ago2* to their target site on the RNA. Mapping the small RNAs from the CLIP-L chimeras to canonical and miRNA hairpin sequences enables capturing mature miRNA and their variants (isomiRs) that interact at the target site (**Fig.** 1B). We adapted the SCRAP pipeline (see *Methods*) to be able to annotate canonical miRNAs and their isomiR types binding to target mRNA sites from CLIP-L chimeras (**Supplementary Fig.** 1).

Across all protocols (**Table ??**), 5′ isomiRs accounted for only ~3% of retrieved miRNA–target interaction pairs (3,393 chimeras), while canonical miRNAs represented ~28% (26,698 pairs) and 3′5′ isomiRs ~11% (11,870 pairs). In contrast, 3′ isomiR–target interactions were the most abundant, comprising ~57% of the dataset (58,911 pairs) (**Supplementary Fig.** 9). Minimum Free Energy (MFE) distributions for both canonical miRNAs and isomiRs over the miRNA–target hybrids revealed dependence on target length: shorter targets produced less stable hybrids (higher MFE), with a steep rise in energy for targets shorter than 20 *nts* (**Supplementary Fig.** 2). To avoid artifacts from these unstable interactions, hybrids with target lengths below this cutoff were excluded from our post-analysis, ensuring that downstream results reflect biologically meaningful and stable interactions.

### 3.2 CLIP-L exhibit sequence bias in miRNA-target chimeras

CLIP-L experiments use linkers that ligate the small RNAs, guided by *Ago* to their target mRNA site. The ligation of the linker sequence is facilitated by a ligation enzyme and the *chimeric read* constituting the miRNA sequence and a part of the RNA target site sequence (**Fig.** 1B). Interestingly, we observed ligation biases at the *first 5 nts* of the 5′ end of the target mRNA. The bias was observed in all protocols and showed reduced preference of the ‘G’ nucleotide at the target site, except for the CLEAR-CLIP protocol where effects were less pronounced (**Supplementary Fig.** 4). In contrast, no prevalent bias on the 3′ ends of the miRNA region could be identified, suggesting no preference by the enzymes based on the miRNA sequence. Sequence logos derived from the target sites further supported these biases, a comparison across protocols (**Supplementary Fig.** 4D) showed that CLEAR-CLIP (Moore et *al*., **Table ??**) was less affected.

MFE distributions of the miRNA and target hybrids from CLIP-L protocols showed energies in the range of unstable hybrids (*>* 5Kcal/mol) (**Supplementary Fig.** 3). Notably, we observed that miRNA target prediction models are affected by the biases present in the data. To our knowledge, such biases have not been reported for CLIP-L data before. Indeed, training a CNN*-baseline* model on the miRNA and target RNA sequence pairs showed inflated performance when the bias was included. A perturbation analysis illustrated a disproportionate reliance on the first four nucleotide positions, an effect inconsistent with biological expectations (**Fig.** 1D). After bias correction, sequence logos displayed a more uniform distribution of the four bases across all positions (**Supplementary Fig.** 10), and the hybrids exhibited MFE distribution towards more stable interactions for both canonical and isomiR interactions((**Supplementary Fig.** 11 A,B), MFE *<* 5 kcal/mol), compared to MFE before bias correction (**Supplementary Fig.** 3A) where the distribution across most protocols had a MFE peak towards 0 kcal/mol, suggesting unstable RNA-RNA hybrids. Thus, correcting these biases prior to model training is essential to ensure that predictions of miRNA–target pairs are driven by true biological signals rather than artifacts.

### 3.3 Ablation studies inform a robust model design

A parameter efficient and robust design for classifying miRNA and isomiR interactions was guided by model ablation studies on three design choices: (a) stacked hybrid attention layers combining self-attention and cross-attention, (b) the bilinear fusion component used for merging miRNA and target features, and (c) the CNN type used in the refinement layers. The hybrid attention reduced variance of performance metrics under cross-validation, making the model more robust to data variability across folds (**Supplementary Fig.** 12A). This held when comparing miRXplain to models with single attention layers (*self-attention (n=1)* and *cross-attention (n=1)*) and to models with two encoder layers of same attention type (*self-attention (n=2)* and *cross-attention (n=2)*) demonstrating the benefit of mixed attention. Replacing bilinear fusion with operations such as either simple concatenation of miRNA and target encoder features or element-wise subtraction led to a significant drop in performance in both cases (**Supplementary Fig.** 12B). Although the performance difference between concatenation and fusion models is modest, the factorized bilinear fusion’s low-rank projection substantially reduces parameters from 1.8 to 1.7M and training time from *>*8 hours to *<*5 hours in 10-fold cross-validation (**Supplementary Table** 4). Finally, among alternative CNN architectures evaluated for the refinement module, the residual CNN achieved the best performance, surpassing basic, depthwise, inception, and dilated CNNs (**Supplementary Fig.** 12C).

### 3.4 miRXplain outperforms state-of-the-art deep learning approaches across miRNA types

To model canonical and isomiR-target interactions from CLIP-L experiments, *miRXplain* combines a hybrid attention mechanism with positional encoding, a low-rank bilinear fusion module, and convolution blocks to refine interaction features (**Fig.** 2). As previously published DL architectures for miRNA target prediction did not use CLIP-L experiments for training, we re-implemented every architecture to create a unified benchmarking setup for systematic performance comparisons (see *Methods*).

In our evaluation, *miRXplain* achieved the best results, reaching in 10-fold cross-validation a mean test auROC and auPRC for canonical miRNA interactions of 0.92 and 0.93, respectively, significantly surpassing CNN*-baseline* (0.84, 0.85) and outperforming recent DL methods, GraphTar (0.80 ± 0.01 SD, 0.81 ± 0.01 SD), DMISO (0.85 ± 0.03 SD, 0.86 ± 0.03 SD), miTAR (0.81 ± 0.01 SD, 0.81 ± 0.01 SD), Mimosa (0.75 ± 0.01 SD, 0.77 ± 0.01 SD), and TEC-miTarget (0.91, 0.92) (**Table** 2). We achieved similar results for isomiRs, with consistent performance improvements across all isomiR types (**Fig.** 3) and over other performance metrics (**Table** 2). Notably, *miRXplain* achieved these results using 15x fewer model parameters than the next best performing model TEC-miTarget, allowing faster training times, lower memory usage, and reduced risk of overfitting, while maintaining superior predictive performance. *Mimosa*, a transformer-based model, underperformed due to its reliance on specific features derived from sequence alignment parameters between RNA sequences that consider only the canonical miRNA form, and which likely do not generalize to the richer CLIP-L data containing isomiR interactions.

**Fig. 3:**
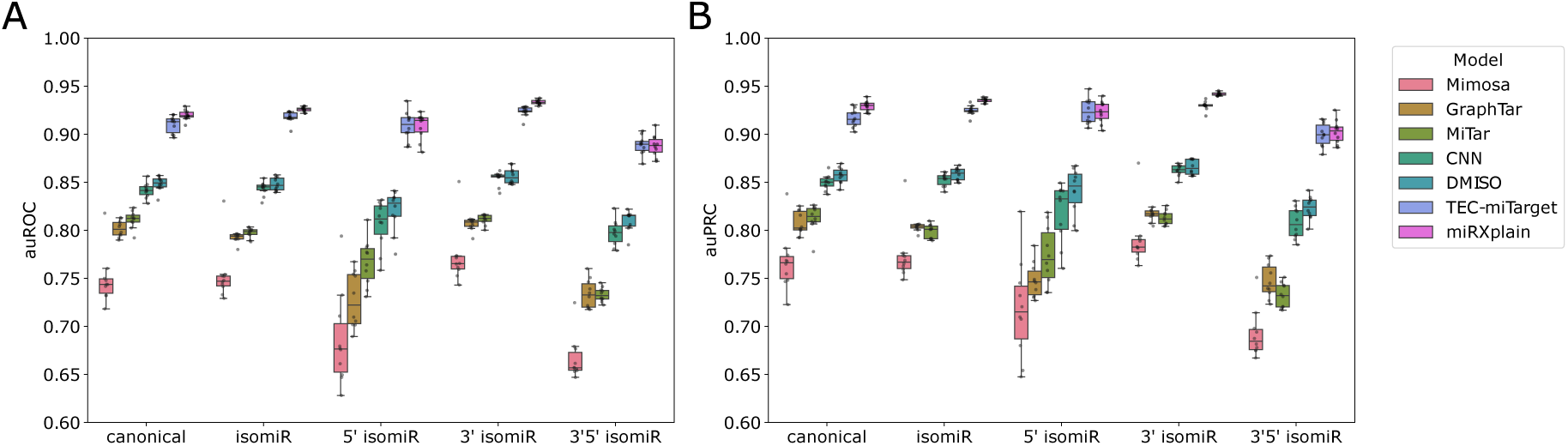
Performance evaluation of recent DL approaches over canonical and isomiR types. Performance metrics **(A)** auROC and **(B)** auPRC of *miRXplain* in comparison to benchmarked models, across miRNA types: canonical, 5′ isomiR, 3′ isomiR, 3′5′ isomiR and isomiR: aggregated for all isomiR types.

**Table 2:** Performance benchmark of *miRXplain* against five most recent DL architectures to predict canonical and isomiR interactions from CLIP-L.

| Model | auROC |  | auPRC |  | Accuracy |  | Precision |  | Recall |  | F1 |  |
| --- | --- | --- | --- | --- | --- | --- | --- | --- | --- | --- | --- | --- |
|  | *can.* | *iso.* | *can.* | *iso.* | *can.* | *iso.* | *can.* | *iso.* | *can.* | *iso.* | *can.* | *iso.* |
| CNN-baseline | 0.84 | 0.84 | 0.85 | 0.85 | 0.76 | 0.76 | 0.75 | 0.75 | 0.80 | 0.80 | 0.77 | 0.78 |
| miTAR | 0.81 | 0.80 | 0.81 | 0.80 | 0.73 | 0.72 | 0.74 | 0.73 | 0.75 | 0.74 | 0.75 | 0.74 |
| DMISO | 0.85 | 0.85 | 0.86 | 0.86 | 0.77 | 0.76 | 0.77 | 0.77 | 0.78 | 0.78 | 0.77 | 0.77 |
| GraphTar | 0.80 | 0.79 | 0.81 | 0.80 | 0.72 | 0.72 | 0.73 | 0.72 | 0.74 | 0.74 | 0.73 | 0.73 |
| Mimosa | 0.75 | 0.75 | 0.77 | 0.77 | 0.68 | 0.68 | 0.68 | 0.69 | 0.71 | 0.71 | 0.69 | 0.70 |
| TEC-miTarget | 0.91 | 0.92 | 0.92 | 0.92 | 0.83 | 0.84 | 0.85 | 0.86 | **0.82** | **0.83** | 0.83 | 0.84 |
| miRXplain | **0.92\*** | **0.93\*** | **0.93\*** | **0.94\*** | **0.84** | **0.85** | **0.87** | **0.87\*** | 0.81 | 0.82 | **0.84** | **0.85** |

*can.*: canonical, *iso.*:isomiR, t-test to test significant differences between *miRXplain* and TEC-miTarget, \* indicates $p < 0.05$ .

**Table 3:** Comparison of model sizes and parameters between the models used in the benchmark and *miRXplain*.

| Model | No. of parameters (M) | Size (MB) |
| --- | --- | --- |
| CNN-baseline | 0.007 | 0.027 |
| miTAR | 0.32 | 1.295 |
| DMISO | 0.11 | 0.444 |
| GraphTar | 0.59 | 2.349 |
| Mimosa | 0.84 | 3.352 |
| TEC-miTarget | 26.7 | 106.767 |
| miRXplain | 1.70 | 6.963 |

### 3.5 Attention mechanism identifies isomiR seed shift effects

Visualizing attention weights can reveal the regions of the input sequence that the model selectively focuses on while learning miRNA interactions. We inspected self-attention maps of the miRNA sequence from the encoder layer of *miRXplain* and observed higher attention weights at specific positions within the miRNA seed region. For isomiRs, the attention weights at the positions were shifted by the number of nucleotides modified on either end of the canonical miRNA. For example *miR-21-5p*, a well-studied oncomiR promoting cell proliferation, migration, and invasion in various cancers, with abundantly expressed isomiRs [58, 59], showed higher attention weights at the 4*th* position (x-axis) in relation to the 2*nd* and 7*th* position (yaxis) of the miRNA seed region (**Fig.** 4A). The model found additional importance at positions 9 and 13 in relation to the miRNA 3′ supplementary regions (11 14 *nt*) (**Fig.** 4A), distinct from other positions. Its isomiR variants (5′ 1nt deletion, 5′ 1*nt* addition, and 5′ 2nt addition) showed shifts of -1 towards the 5′ end, +1 or 1*nt* towards the 3′ end, and +2 or 2*nt* towards the 3′ end, respectively (**Fig.** 4 B-D) relative to its canonical form. Attention maps were background corrected to remove spurious artifacts (**Supplementary Fig.** 13). The analysis suggests that nucleotide modifications in miRNAs could induce changes in target site selection.

**Fig. 4:**
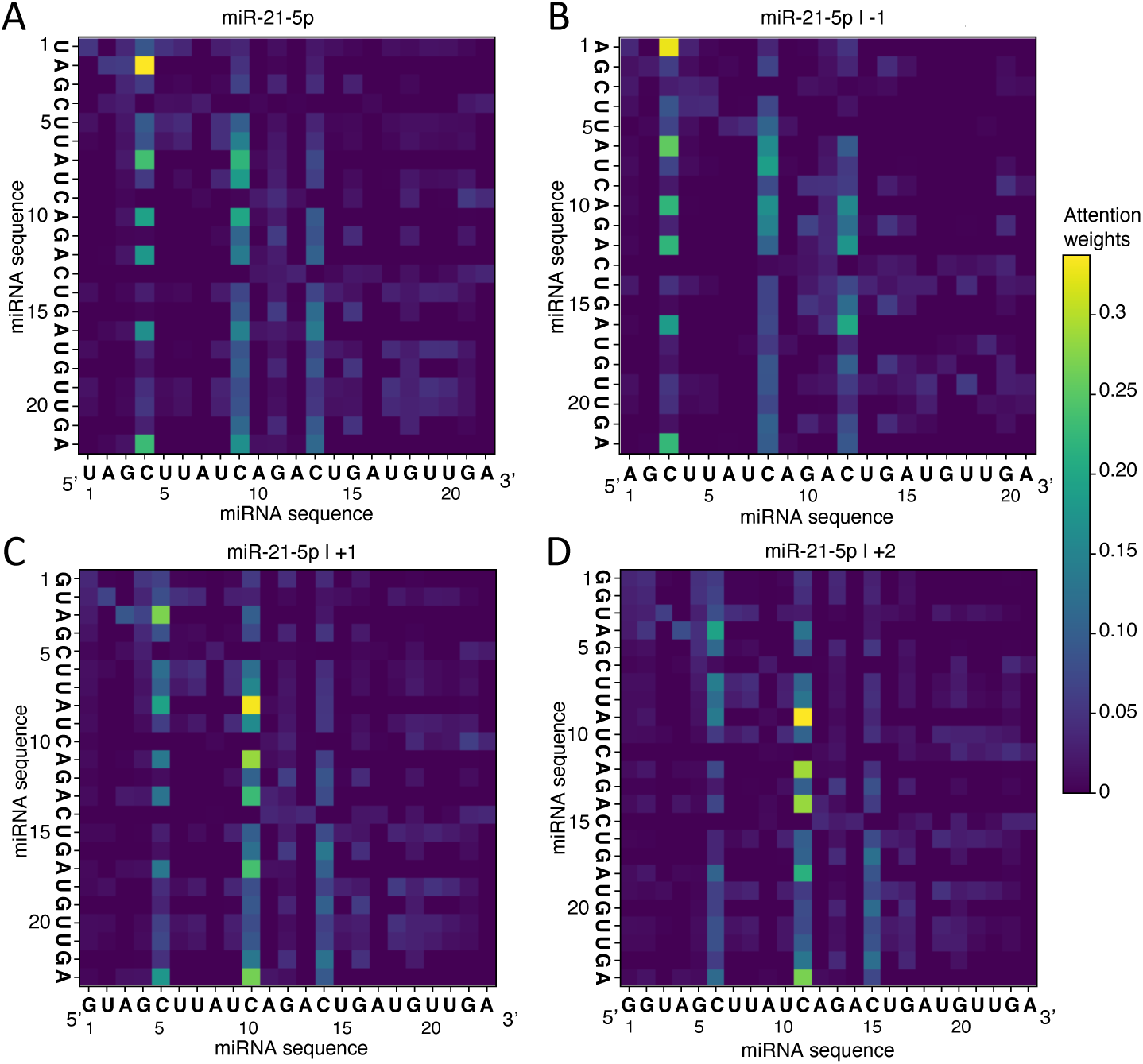
Self attention maps highlight differences in canonical and isomiR interactions with their targets. Self-attention maps of miR-21-5p and its isomiR variants from *miRXplain* reveal differences in model attention weights. Shown are the canonical form **(A)**, isomiR variations on the 5′ end with 1 *nt* deletion **(B)**, 1 *nt* addition **(C)**, and 2 *nt* addition **(D)**. The positions of the miRNA sequences are shown below the respective nucleotides. Columns with high attention weights shift depending on the isomiR event, reflecting a change in binding importance.

### 3.6 In silico saturation mutagenesis reveals importance of miRNA seed and 3*’* supplementary regions

*In silico* saturation mutagenesis (ISM) was employed to reveal the importance of the nucleotide position from the interacting RNAs as measured by the prediction probability changes upon mutations. We visualized overall *in silico* mutation effects by averaging prediction deltas Δ*p* aggregated over different miRNA/isomiR types in binding at each nucleotide position over the miRNA/isomiR sequence or the target sequence of the CLIP-L chimera. We observed higher magnitudes of mutation effects in the miRNA seed (position 2-8) and the 3′ supplementary regions (pos 13-16), for canonical miRNAs and isomiRs (**Fig.** 5), thereby recapitulating important miRNA regions in miRNA target interactions.

**Fig. 5:**
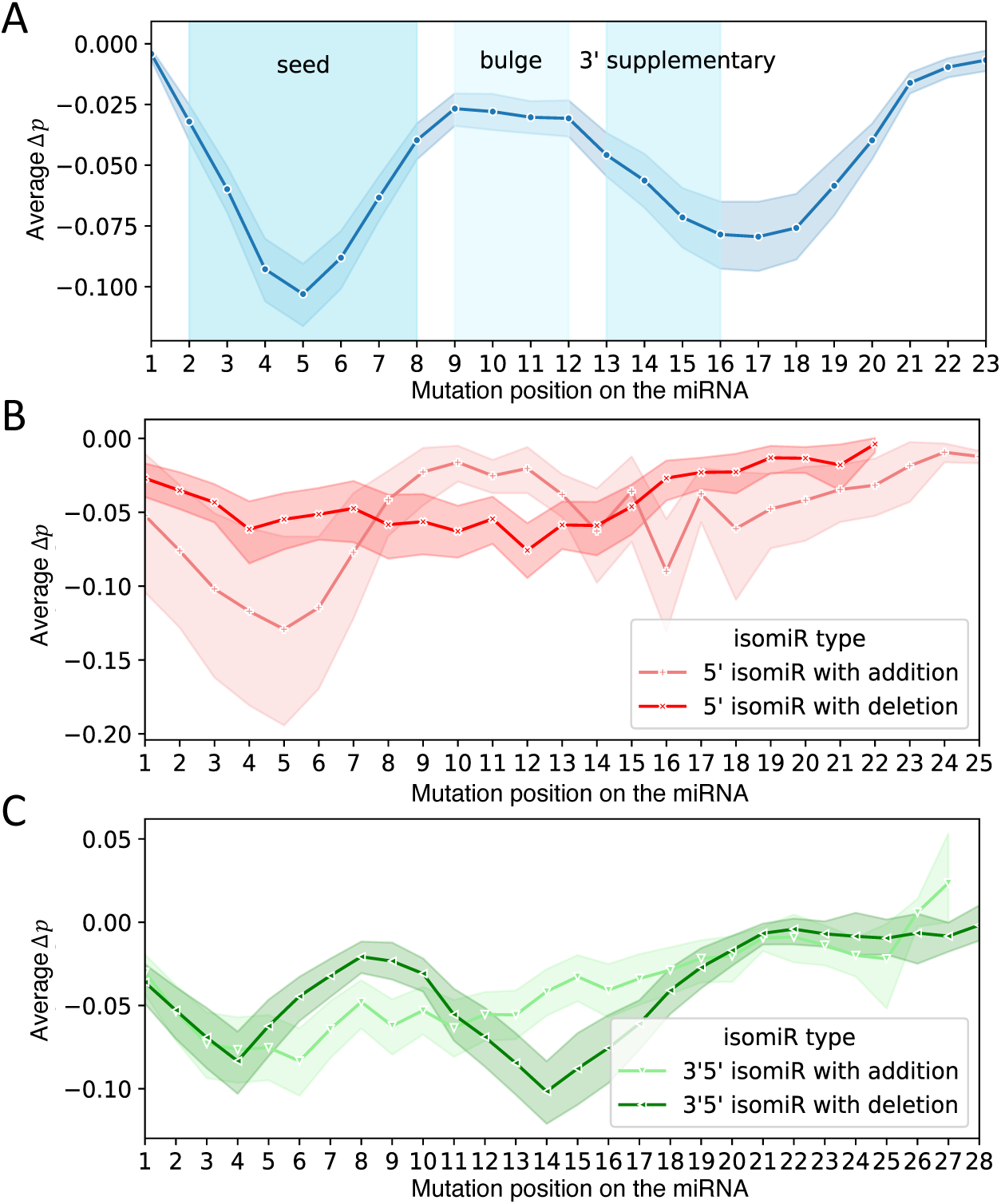
*In silico* mutagenesis analysis identifies miRNA seed, bulge and 3′ supplementary regions. **(A)** Analysis of canonical miRNAs with sequence length up to 23 *nts*. Average prediction delta values (Δ*p*) on the Y *axis* are shown as a function of the mutation position introduced along the miRNA (X *axis*). The shaded areas highlight the seed, the bulge, and 3′ supplementary regions. **(B-C)** Delta values for isomiR types with addition or deletion of nucleotides for **(B)** 5′ isomiRs, and **(C)** 3′5′ isomiRs. For addition and deletion events, 5′ isomiRs have a majority of +1 *nts* and -1 *nts*. For 3′5′ isomiRs, we annotate ‘addition’ or ‘deletion’ events as modifications at the 5′ ends. 3′5′ isomiRs with addition events have a majority of 2 *nts* at 5′ end and -4 *nt* at 3′ end, while deletion events have a majority of -1 *nt* at 5′ end and -1 *nt* at 3′ end.

To investigate if our model captures the effects of isomiR variations, we split isomiR types into variants with additions or deletions of nucleotides at the ends. We identified *seed shifts* towards the 5′ or 3′ end of the sequence in case of modifications at the 5′ end of the miRNA, which applies to both 5′ and 3′5′ isomiRs (**Fig.** 5). In contrast, modifications at the 3′ end of 3′ isomiRs did not impact the *average* Δ*p* profile, following a similar profile to that of the canonical miRNA (**Supplementary Fig.** 14). This agrees with known reports that modifications at the 5′ ends of the mature miRNA exert *seed shift* effects on target selection [7]. We find more pronounced effects in the beginning, towards the 5′ end of the CLIP-L target, as most of the miRNA/isomiR binding is in the beginning of the target site [23]. Higher Δ*p* was observed until the 10 12 *nt*, following which it drops until the length of the miRNA target (**Supplementary Fig.** 15). Moreover, as there is no ground truth measurement of the exact position of miRNA binding on the target, it was not possible to positionally align the target sites, compared to region-based annotation for miRNAs. ISM analysis on the target region from the CLIP-L chimera did not reveal any distinguishing regions as effect, measured as changes in Δ*p*) among miRNA/isomiR types.

### 3.7 miRXplain identifies canonical and isomiR targets and associated functions

In our analyses thus far, we used *miRXplain* in the context of individual binding sites. As there exist several databases that list experimentally validated canonical miRNA–gene interactions, these can be leveraged to evaluate machine learning models. In contrast, no comparable resources exist for isomiRs. As a result, *miRXplain* cannot be benchmarked at gene-level resolution for isomiRs specifically. Therefore, we sought to investigate the differences between canonical miRNA and their respective isomiR forms in gene-level analyses relying on reported canonical target genes.

We designed a statistical model that combined predicted binding events of *miRXplain* in nonoverlapping bins of a gene’s 3′ UTR (**Fig.** 6A, see Methods). For example, when applying our strategy to miR-21-5p, a known oncomiR, we observed that experimentally validated target genes [54, 55] had significantly larger gene *p* values (-log10) than non-targets (Wilcoxon rank sum test, p*<*0.05, **Fig.** 6B). This supports the validity of *miRXplain* and the statistical aggregration of individual binding sites on gene level.

**Fig. 6:**
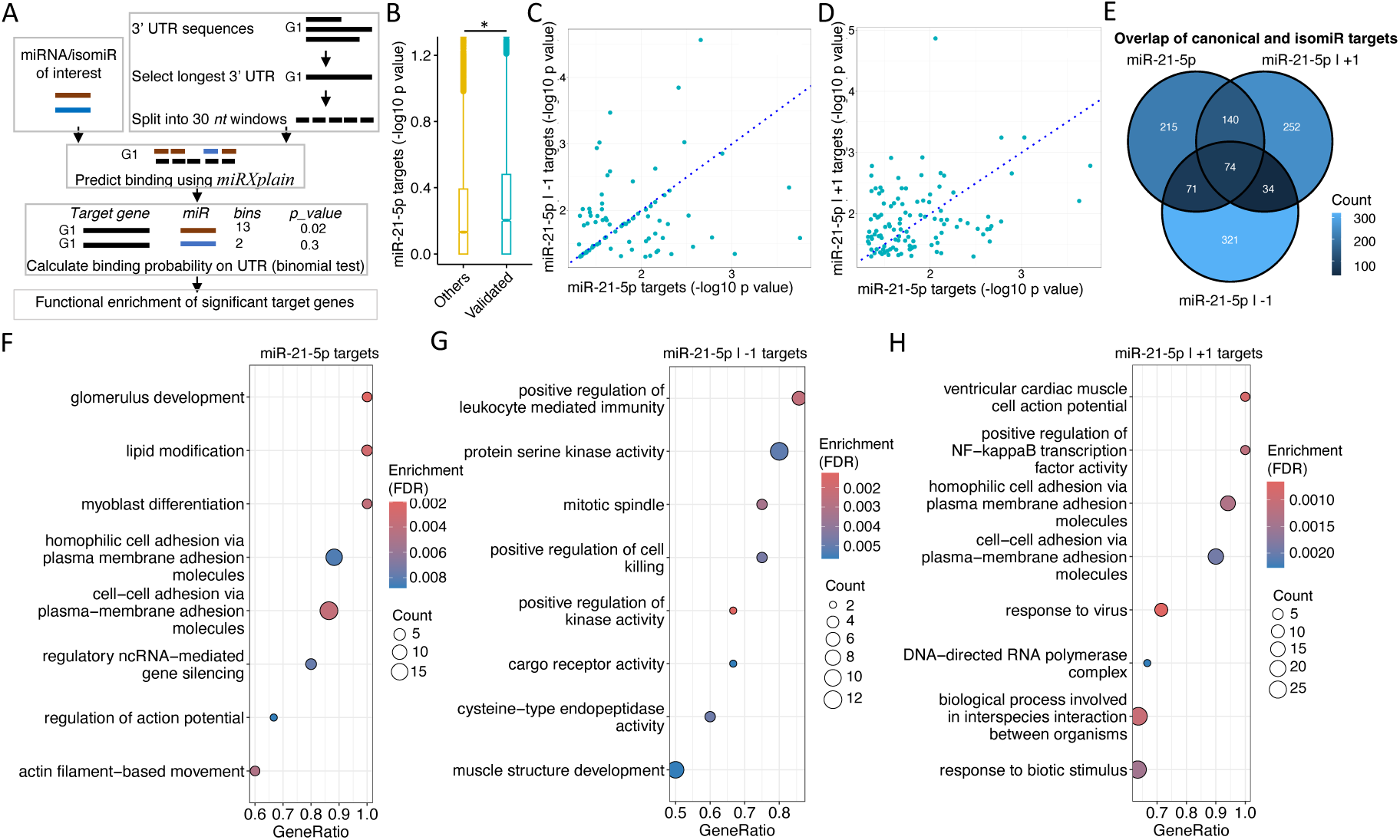
*miRXplain* highlights canonical and isomiR targets and associated functions. **(A)** Overall workflow to predict miRNA and isomiR target genes using *miRXplain*. **(B)** Analysis of experimentally validated miR-21-5p target genes (Validated) against genes for which no regulation of miR-21-5p is known (Others). Gene *p* values (-log10, Y *axis*) obtained using *miRXplain* are contrasted between the two gene sets. Difference in gene ranks was assessed using Wilcoxon rank sum test (‘*’ *p <* 0.05). **(C, D)** Analysis of differences in canonical and isomiR binding on validated target genes. Gene *p*-values are compared between canonical miR-21-5p (X *axis*) and miR-21-5p isomiRs (Y *axis*), with -1 deletion **(C)** and +1 addition events **(D)**. Each dot denotes one target gene. **(E)** Overlap of top 500 canonical and isomiR targets of miR-21-5p. **(F-H)** Gene set functional enrichment results, in decreasing order of gene ratio, for top 500 target genes of canonical miR-21-5p **(F)**, -1 **(G)**, and +1 isomiR targets **(H)**. The X *axis* shows the gene ratio, the amount of genes annotated with a certain function (Y *axis*). Each circle is colored to represent the significance of the associated functions (FDR Hypergeometric test), with the size proportional to the count of genes annotated.

Interestingly, two isomiRs of miR-21-5p have previously been investigated for gene regulation [60]. We compared the gene *p* values for canonical miR-21-5p and its isomiR miR-21-5p -1 on the known canonical target genes (**Fig.** 6C). We found that for a large fraction of target genes, there was no difference in gene *p* values, and when there were differences, no clear trend was apparent. However, comparison to isomiR mir-21-5p +1 on the known target genes showed much stronger differences compared to the canonical form (**Fig.** 6D). When comparing the top *predicted* 500 genes for the canonical and the two isomiR forms of miR-21-5p, we found that only 14,8% overlapped between all three miRNAs (**Fig.** 6E). As there exists no validated isomiR targets, it remains uncertain how different canonical targets are from their isomiR forms.

The top 500 targets identified for the canonical form of miR-21-5p showed enrichment for many previously reported functions of this miRNA, such as: *glomerulus development* [61], *protein serine kinase activity* [62] and *cell-cell adhesion* [63] in prostate cancer, *myoblast differentiation* [64] and *mitotic spindle* regulation [65](**Fig.** 6F). We also conducted enrichment analysis for the two isomiR variants (**Fig.** 6G, H). The most enriched terms differed from the analysis of the canonical targets, which is to be expected as the genes varied substantially (**Fig.** 6E). Noteworthy, mir-21-5p +1 genes where enriched for regulation of NF-kappa*ω* pathway activity, which was reported to be regulated by the canonical form of miR-21 [66, 67].

As a comparison, we conducted a similar analysis for predicting target genes of miR-9-5p and its known isomiR miR-9-5p -1. Again we found that *miRXplain* gene *p* values of experimentally validated target genes of miR-9-5p were significantly smaller than non-target genes (Wilcoxon rank sum test, *p <*0.05)(**Supplementary Fig.** 17A). Also there was little similarity between gene *p* values of the isomiR and canonical form and although the overlap between target genes is substantial (42% shared targets among the top 500) the majority differed (**Supplementary Fig.** 17B,C). Gene enrichment analysis of the canonical form for miR-9-5p revealed terms that are supported by recent literature: *inhibition of interleukin-6* [68], the *JAK/STAT3 pathway* [69], and *regulation of endothelial cell migration and angiogenesis* [69, 70] (**Supplementary Fig.** 17D). However, the isomiR miR-9-5p -1 showed enrichment of other functional categories such as target genes related to the *polymerase 2 complex* and *response to tumor necrosis factor* (**Supplementary Fig.** 17E).

Our analysis of target genes of two miRNAs and their isomiRs suggests that isomiRs enable regulation of a potentially diverse set of genes, compared to the canonical form, that can drastically differ in their function. Previous analyses of isomiR and canonical targets had also found differences in the involved gene functions [71, 72], which underlines the importance of studying the role of individual isomiRs in more detail.

### 3.8 miRXplain captures SNV effects

Sequence-based deep learning models are widely used to prioritize functional variants in non-coding regions and RNA regulatory elements by evaluating whether a point mutation significantly alters the model’s prediction probability—for example miRNA-target binding probability. *miRXplain* allows investigation of the effect of Single Nucleotide Variants (SNVs) that associate with clinical phenotypes by inspecting the predicted probability of miRNA–target binding for reference sequences and after introducing a mutation.

In order to carry out such an analysis, we first mined a large set of GWAS conducted on UK Biobank data and intersected the location of significant SNVs, linked to different phenotypes, with annotated canonical miRNAs (see Methods). We found 28 SNVs that overlapped with 6 canonical mature miRNA coordinates which had interactions in the CLIP-L experiments (miR-589-3p, miR-450a-5p, miR-30a-3p, miR-1910-5p, miR-1224-3p, and miR-3117-3p) (**Supplementary File** 1).

Second, we used *miRXplain* to compute the changes in the predicted interaction probability caused by the SNV. With an analysis similar to that of the previous section (**Fig.** 6A), we calculated the binding probabilities for the reference miRNA sequence and with the SNV allele introduced, comparing binding probability on the 3′ UTRs of all validated targets, using the binomial statistic (Eq. 21, see Methods). For 5 out of the 6 miRNAs a significant variation in binding *p* values was observed (Wilcoxon rank sum test, p*<*0.1). For example, the SNV (rs372099307) overlapped miR-589-3p in the 3′ supplementary region close to the tail at position 20*nt* (**Fig.** 7A, C). The importance of the 3′ supplementary region for miRNA target identification was recently reported [73]. The mutated miR-589-3p showed diverse, but often reduced binding probabilities compared to its canonical form (**Fig.** 7A). Another SNV (rs9271448) overlapped miR-30a-3p at position 9*nt*, with a similar pattern (**Fig.** 7B). The differences for other SNVs overlapping miRNAs are shown in **Supplementary Fig.** 18, except miR-1224-3p, for which the differences were not significant. This analysis suggests that GWAS SNVs overlapping miRNAs have the potential to significantly change miRNA binding behaviour in validated target genes.

**Fig. 7:**
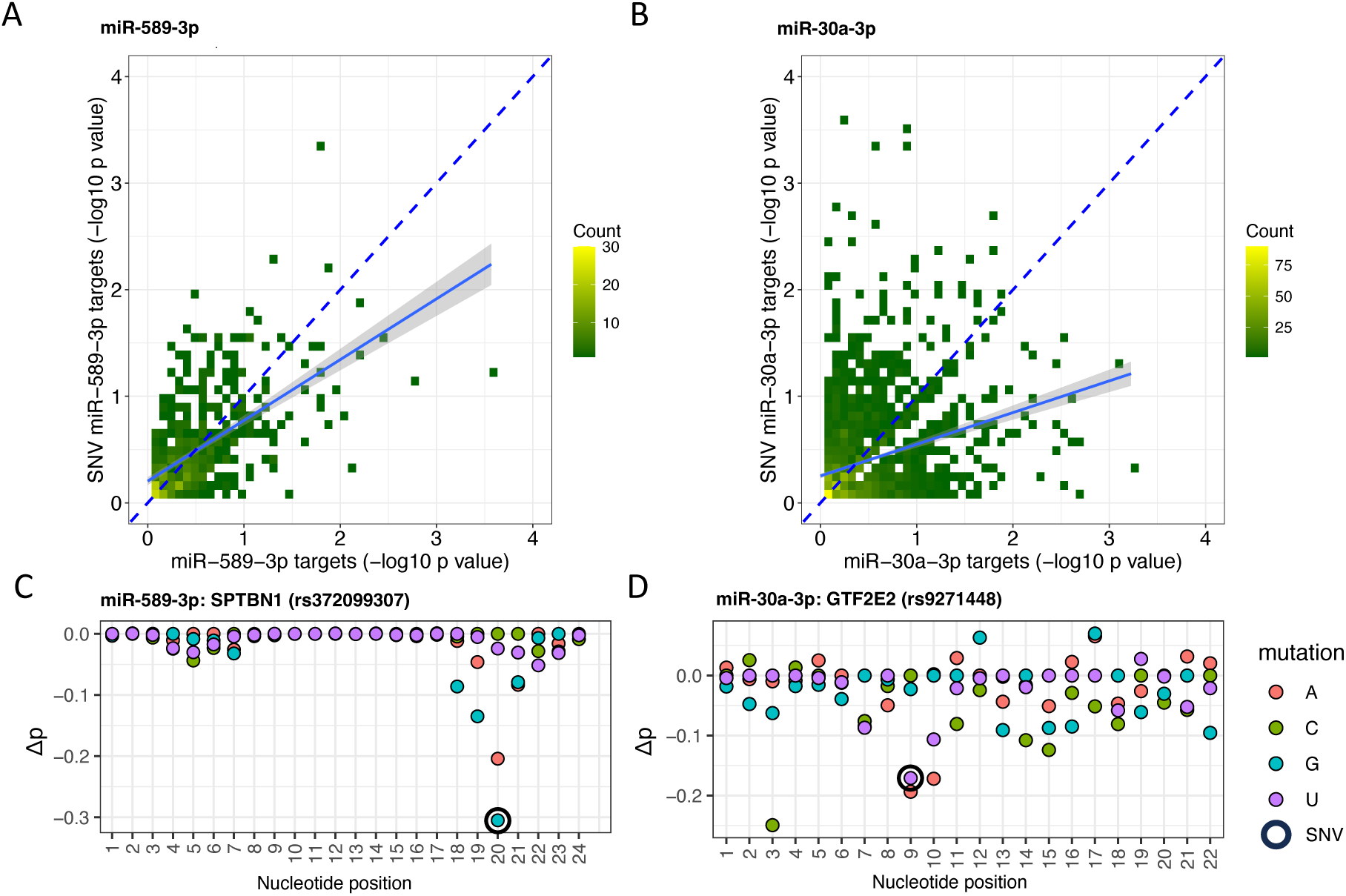
Predicting SNV effects in mature miRNAs using miRXplain. *miRXplain* was used to investigate the effects of an SNV in miRNAs. Target gene binomial *p* values (workflow Fig. 6A) were computed for the miRNA with the SNV (Y *axis*) and the reference sequence (X *axis*). The binned scatter plots show the distribution of the *p* values (-log10) for all validated target genes for miR-589-3p and SNV rs372099307 **(A)** and miR-30a-3p with SNV rs9271448 **(B)**. The regression line (in solid blue) indicates a deviation from the diagonal (in dashed blue). Differences between both between the canonical and the mutated miRNA forms were significant (Wilcoxon test p *<* 0.1). Detailed analysis of binding probability with miR-589-3p on a SPTBN1 binding site **(C)**, and miR-30a-3p on a GTF2E2 binding site **(D)**. *In silico* mutation results in a decrease in Δ*p* if the binding is unfavorable or an increase if binding is favorable (Y *axis*), resulting in negative or positive values at the SNV position (X *axis*). The reported SNV on the miRNA is encircled (in black). *Note*: As both miRNAs are encoded on the negative strand, reference and SNV alleles are shown as reverse complements.

Another way to investigate the effect of SNVs on miRNA binding, is to study individual binding sites using *miRXplain*. Analysis of an interaction of miR-589-3p with the SPTBN1 (*ω* II-spectrin) gene is illustrated in **Fig.** 7C. SPTBN1 is a molecular scaffolding protein with functions in cell shape determination, transmembrane protein arrangement, and organelle organization and mutations in SPTBN1 lead to a neurodevelopmental syndrome [74]. The SNV is associated with benign neoplasm of thyroid gland (ICD-10: D34). miR-589-3p with the observed SNV showed the strongest change in binding probability among all possible mutations for that binding site.

Using a similar analysis, a reported SNV for miR-30a-3p was found at the 9th *nt* in the bulge region (**Fig.** 7D). The SNV in miR-30a-3p is associated with psoriasis in male (20002 1453). Mutation in the bulge region resulted in the fourth largest decrease in prediction probability observed for that miRNA binding site on the GTF2E2 gene. Note that the deviation in Δ*p* values for the SNV mutation is much larger compared to the average profiles (**Fig.** 5). GTF2E2 (general transcription factor IIE) is part of the RNA polymerase II transcription initiation complex, an essential ubiquitously expressed gene in many tissues. Mutations in GTF2E2 are known to cause trichothiodystrophy [75], a disease which may come with skin issues for patients. These analyses illustrate how *miRXplain* can be used to study the effect of SNVs in a miRNA across many target genes or for studying effects on individual binding sites.

## 4 Discussion

### 4.1 CLIP-L is superior to CLIP in capturing miRNA binding specificities, but contains biases

#### Choosing CLIP-L experiments over CLIP

One of the main contributions of this work is that *miRXplain*, unlike previously developed deep learning tools for predicting mRNA–miRNA interactions, leverages the full breadth of ligation-based CLIP-L protocol data only, providing a theoretical and practically experimental advantage over existing approaches. Apart from the incorporation of the Helwak Ago CLASH dataset in *DMISO* and a few others, current methods rely predominantly on experimentally derived miRNA–target interactions from curated resources such as DIANA TarBase [55], miRTarBase [54], or as done in mirMark [76], which are mostly built from Ago PAR-CLIP and HITS-CLIP datasets. These protocols are non-ligation-based and therefore do not preserve direct miRNA–mRNA chimeric information. Because CLIP, without ligation step, does not identify the exact miRNA that guides Ago to their putative target, early deep-learning–based miRNA target prediction methods trained on these datasets were needed to refine positive interactions using heuristic rules—such as seed complementarity, hybridization energy, or additional sequence features. This strategy was partly driven by the fact that the CLIP dataset is large enough to support training of deep learning methods and because deep neural networks are generally robust to noise. However, this reliance on indirect evidence ultimately limits the precision with which models can learn the true miRNA–mRNA pairing rules; the assigned miRNA might not correspond to the small RNA actually bound [38]. In addition, as they cannot capture exact miRNA variants, models trained mostly on CLIP data offer limited interpretability for discovering novel miRNA biology, in particular when it comes to isomiR target interactions.

By contrast, *miRXplain* exploits the richer pairing information captured by ligation-based CLIP-L protocols, allowing the model to learn interaction features from true chimeric reads rather than inferred miRNA-target pairs, and allows transcriptome-wide analysis of both canonical miRNA as well as isomiR-target interactions (**Fig.** 1B). As we show, this leads to improvements in predictive performance for all miRNA and isomiR classes, and represents a meaningful step forward in the development of miRNA target prediction tools grounded in experimentally direct evidence.

#### CLIP-L experiments contain sequence biases in the target region of chimeric reads, which must be corrected before modeling

Although CLIP-L is specific in capturing exact miRNA–target pairs and thus offers a clear advantage over non-ligation-based protocols, it is affected by protocol-induced biases. Our analysis revealed sequence biases within the first four nucleotide positions of the target site (**Fig.** 1C, **Supplementary Fig.** 4), most likely due to ligation enzymes that tether the miRNA/isomiR with the target site. RNAse T1 treatments have been shown to have pronounced G-depletion in the binding sites, as used in CLIP-L experiments, especially during the ligation steps [77, 78]. Another possible influence is the choice of sRNA library construction protocol, as these methods can introduce characteristic nucleotide biases [79]. We show that using such biased target sequences for modeling miRNA-target interactions inflates model performance and distorts interpretation (**Fig.** 1D). We thus processed and corrected the target sites to account for this experimental bias.

To remove the identified bias, we trimmed 4 nucleotides from the 5′ end of all target sequences, thereby removing bias-affected positions. Subsequently, to ensure that the trimmed region was not involved in binding and to report the precise miRNA–target contact points, we assigned binding locations using IntaRNA [44], a method that computes the miRNA and target binding positions, while accounting for duplex hybridization energy, target accessibility, RNA secondary structure, and seed constraints, and ranks among the top methods for sensitivity and precision across independent benchmarks [80]. After running IntaRNA, we filtered out all interactions where miRNAs or their variants were predicted to bind within the bias-affected trimmed nucleotide positions (**Fig.** 1C).

Correcting for protocol-induced biases in our datasets proved essential. This was illustrated by experiments with a simple CNN model, where training on full target sequences—containing the characteristic bias in the first 4 nt—led to artificially inflated performance compared to training on trimmed sequences that mitigate this bias. The model trained on uncorrected data primarily learned the protocol-specific sequence artifact rather than the true biological determinants of miRNA–target interactions. These results underscore that the success of deep learning methods depends not only on leveraging large and diverse training datasets but also on careful data curation to minimize systematic biases that can mislead the model and compromise its generalizability.

As novel high-throughput protocols for the identification of miRNA/isomiR–target interactions and improved ligation strategies are being developed, the increasing quality of available data will enable the training of more reliable computational models [81]. Ideally, bias-free CLIP-L protocols would allow these datasets to be used directly, without the need for ad-hoc post-processing steps such as deciding how to remove protocol-induced biases or filtering interactions after bias correction, as we were required to do here. Such steps inevitably introduce arbitrary choices and reduce the size of the training dataset, potentially limiting the model’s performance. Eliminating protocol bias at the experimental level would therefore greatly improve data quality while preserving the full breadth of the dataset for model training.

### 4.2 Transformer-based sequence models learn miRNA target binding specificity

#### Choice of miRXplain architecture for classification and interpretation

Among recent deep-learning approaches, many methods have relied on pre-processed CLIP datasets that already encode assumptions about canonical seed pairing. In several cases, the training sets were partially pre-filtered using seed-complementarity rules, as in miTAR [33], or placed much emphasis on specific miRNA regions (e.g., the 5′ seed region [34], thermodynamic constraints, or sequence conservation [35]). Such heuristics risk creating a circular evaluation scenario in which models perform well primarily on datasets constructed using the same rules embedded in the positive training set. For instance, miTAR identified the miRNA seed as the dominant predictive feature—an unsurprising outcome given that its training data had already been filtered by seed complementarity. Such predefined constraints may limit a model’s ability to generalize to ligation-based CLIP-L datasets, where interaction logic is more complex.

DMISO [36] presented an early effort to learn isomiR–target interactions from CLASH and CLEAR-CLIP data using a hybrid CNN–LSTM architecture. However, the method did not analyze in-depth its input data generated with ligation-based protocols and therefore could not detect or correct nucleotide composition biases that might arise in such libraries. It also did not explicitly model isomiR heterogeneity or attempt to interpret isomiR biology using dedicated interpretation methods. Most critically, the full pipeline and code used to train and evaluate the model were not made publicly available, preventing its use for this task. TEC-miTarget [34] was the first Transformer model developed for the miRNA-target classification task. Although it outperformed state-of-the-art models based on hybrid CNN-RNN architectures, it represented a step back with respect to DMISO by leveraging previously compiled CLIP-derived datasets, therefore not addressing isomiR-specific predictions.

*miRXplain* was designed to overcome the limitations shared by these state-of-the-art approaches. Its transformer-based architecture integrates positional encoding, hybrid self- and cross-attention mechanisms, and bilinear fusion layers to model direct dependencies between miRNA, including isomiRs, and target nucleotides (**Fig.** 2). By avoiding reliance on seed-based heuristics and by learning interaction patterns directly from sequence–sequence relationships, *miRXplain* can capture both canonical and noncanonical binding modes characteristic of CLIP-L interactions. Despite being trained from scratch on only a hundred thousand interaction pairs, the model’s compact parameterization—exemplified by the bilinear fusion component that combines lower-rank representations of the miRNA and its target (**Table ??**)—helps prevent overfitting and allows effective learning from experimentally grounded datasets without unnecessarily increasing model complexity. This likely contributes to *miRXplain*’s superior performance: its architecture is expressive enough to model fine-grained miRNA–target dependencies, yet parameter-efficient enough to generalize beyond the biases of processed CLIP-derived datasets. Consequently, *miRXplain* yields interpretable predictions more consistent with known—and emerging—miRNA biology.

#### IsomiR and canonical interactions are predicted with similar accuracy

While there are many studies that highlight the abundance of isomiRs in different cancer types or tissues [82, 83], there are few dedicated methods for predicting isomiR target sites and no databases with reported isomiR targets. *DMISO* is the only method that explicitly mentions isomiRs, but as detailed above, its software is not available for use. Other studies that investigated isomiR targets relied on existing methods, such as TargetScan, that were not developed for isomiRs [84, 85]. It is currently unclear how well tools designed for canonical targeting perform for isomiRs. Our evaluation across different isomiR classes showed that the investigated deep learning methods perform similarly on isomiRs and canonical miRNAs, with differences between isomiR classes that can be attributed to the varying number of interactions per class. (**Fig.** 3). But as isomiR interactions were part of the training data, a similar performance can be expected. Thus, additional analysis using CLIP-L derived isomiR interactions and other established tools are needed in the future.

#### Interpreting binding finesse of miRNAs and their variants

We interpreted the model to investigate whether canonical miRNAs and their isomiR variants exhibit distinct binding patterns as inferred from nucleotide-level importance. Despite lacking prior knowledge on miRNA type or binding rules, post-hoc interpretation via *in silico* saturation mutagenesis identified differentiating features in binding specificity between canonical and isomiR forms within annotated miRNA regions (**Fig.** 5).

Focusing on canonical miRNAs (**Fig.** 5A), we observe the largest changes in prediction probability along the seed region (nucleotides 2-8), indicating its importance compared to other locations in the sequence. In contrast, for isomiR types with modification at their 5′ end (5′ isomiRs and 3′5′ isomiRs) we report *seed shifts* when compared to the canonical annotated regions and to the position of maximum probability drop on average (position 5 for canonicals). Shifts can be better visualized when differentiating between isomiRs with addition, and deletion events, with the direction of the shift depending on the modification type (**Fig.** 5**B,C**). Differently, 3′ isomiRs (**Fig.** 14) exhibit no seed shifts and their ISM profiles reproduce with high similarity that of the canonicals.

Beyond the seed, the adjacent region—referred to as *bulge* or *central region* (nucleotides 9-12)—is generally excluded from direct binding to the target because less favored for contiguous pairing and is instead structurally occupied by *Ago2* proteins [1]. Although evidence for a consistent targeting role of this region is limited, one study [86] reported that the nucleotides at this region might concur to binding upon rearrangement or extension of the seed, potentially enhancing target repression. Our model interpretation confirms that the involvement of the bulge in miRNA targeting is limited and not consistently observed in our data, and assigns to this region low average importance.

In contrast, the 3′ supplementary region (nucleotides 13-16) highlights positions relevant for binding, consistent with studies that this region improves site affinity and efficiency [87, 88]. Its importance in finetuning miRNA targeting specificity has also been reported in the context of isomiRs with modifications at the 3′end, where increased tail length has been hypothesized to increase stability of supplementary interactions [89].

Taken together, these results indicate that *miRXplain* successfully captures target binding variation across miRNA types, and identifies the sequence positions most crucial for binding. Because RNA structural accessibility influences target binding and hence repression, integrating structural modalities such as *in-vivo* mRNA target site structural profiles obtained through protocols like icSHAPE [90] or DMS-Seq [91] would be beneficial for future model development and interpretation. In particular, high-resolution structural studies, including NMR or cryo-EM, could confirm the differences in binding specificity and conformational variation between isomiRs and canonical miRNAs at their target sites.

#### Target context indicates the presence of additional regulatory regions

Alhough this study focused on the binding specificity of miRNAs and their variants with their CLIP-L targets, we examined the model’s prediction probabilities for increasing target lengths. A minor increase in accuracy was found when extending target sites to 25*nts* at the 3′ ends (**Supplementary Fig.** 6), suggesting the presence of possible regulatory sequence regions or signals in the miRNA-binding neighborhood that might aid hybridization [92]. This is in accordance with the study by Kim *et al*., which reported enrichment of regulatory regions and RBPs upto 50 bps upstream and downstream of miRNA binding sites that aid accessibility for miRNA binding [92].

### 4.3 Evaluation of mutations in miRNA regulation

SNVs on miRNAs can disrupt or promote miRNA target binding, leading to gain or loss of gene regulatory functions. Using SNV-mutated miRNAs, our model revealed miRNA effects associated with disease allele variations. As highly expressed miRNAs or genes can have a high relevance for the cell, our case study considered well-known miRNAs, such as miR-30, reported with many regulatory functions [93, 94], miR-589 is expressed in breast cancer and reported with aberrant expression in various other cancer types [95]. A mutation within the miRNA gene or its processed mature miRNA form can have a significant impact on downstream target binding and affect target expression or translation. Based on what *miRXplain* learnt about the rules of miRNA-target binding, we demonstrate changes in predictions for SNV-associated nucleotides with human phenotypes (**Fig.** 7). However, a comprehensive statistical assessment of whether disease-associated SNVs significantly affect miRNA–target binding is not feasible in this setting. Our analysis is restricted to SNVs occurring within CLIP-L–captured miRNAs and their targets, and the number of such variants (approx 30) is far too small to support a robust, genome-wide evaluation.

### 4.4 Limitations of this study

Different seed binding patterns, especially in relation to canonical and non-canonical miRNAs, with or without binding at the 3′ supplementary region have been reported to affect target repression [87]. Currently, there is a lack of experimentally verifiable negative miRNA target sites, which limits model interpretability. Our current assumption is that the predicted MFE between miRNA and target site provides a good measure for selecting negative miRNA interactions that are not completely trivial, such as random RNA sequences. However, if such negative sites could be experimentally obtained for each miRNA, this could improve the generalization of miRNA-target prediction models.

The preference of CLIP and CLIP-L based protocols for G-poor 3′UTR sites, identified as sequence bias in our dataset, limits our ability to study rules specific to such sites. In this study, we removed the detected nucleotide biases in CLIP-L target RNAs by trimming the biased bases before training. Other types of biases of more complex nature might exist in CLIP-L, such as those due to RNA structure stability or miRNA/isomiR preference for ligation to targets, and could further influence model performance assessment. The development of bias-agnostic experimental protocols to query miRNA target preferences would be crucial for model training in the future.

Our model interpretations are limited in that they overlook dependencies on external regulators that may support miRNA interactions, such as RBPs in the miRNA-target binding neighborhood. They may also be limited by the smaller number of miRNA site interaction instances in the CLIP-L data used for training. The number of miRNA target chimeras in the experiment depends on miRNA abundance and its target gene expression, and thus influences which types of interactions will be learned by models from the CLIP-L data we used.

In addition, miRNA target binding is not the only necessary condition for target transcript repression and miRNA/isomiR mediated functional regulation [96, 97]. Additional factors include miRNA concentration and stoichiometric effects, competition between miRNAs for target selection, transcriptional alterations that may change binding sites accessibility, and binding affinity to CDS and 5′ UTR regions of the mRNA [5]. Our approach brings us one step closer to meaningful identification of miRNA/isomiR and its targets.

## 5 Conclusion

miRNAs are key regulators of diverse biological processes and understanding the rules governing miRNA and isomiR target binding is essential for accurate target selection. We developed *miRXplain*, a deep learning model with the power to reveal and distinguish the preferential target selection of canonical miRNAs and isomiRs.

By addressing ligation-induced nucleotide biases, we curated a CLIP-L derived training dataset for model development aimed at decoding miRNA- and isomiR-specific targeting. Future benchmarking of deep learning and statistical approaches for miRNA target prediction should rely on CLIP-L data annotated with the specific miRNA or isomiR variant involved in the interaction. As a practical resource for the miRNA biology community, *miRXplain* supports binding site-level prediction of miRNA–target affinities and, most critically, facilitates genome-wide interrogation of isomiR targeting and disease-associated SNV effects. *miRXplain* positions itself at the frontier of computational methods advancing the underexplored domain of isomiR biology, paving the way for future efforts to dissect isomiR-specific functional and regulatory mechanisms.

## 7 Code and data availability

*miRXplain* is implemented in Python 3.11, pipelines to preprocess training data are implemented in bash and Snakemake, and associated visualizations are produced with Python and R. All scripts and data used to train the model are publicly available on Github (https://github.com/marsico-lab/mirxplain) and Zenodo (https://doi.org/10.5281/zenodo.18010234).

## 9 Acknowledgments

We acknowledge Florian Neukamm, Goethe University, Frankfurt am Main, for the maintenance and setup of computational infrastructure. We thank Nina Baumgarten for discussion about SNV analysis in the UK Biobank.

## 10 Author contributions

R.K.M. study conception, training data processing and analysis, SNV and miRNA target analysis, writing; G.C. model design, training data processing and analysis, benchmarking, model interpretation, writing; H.C. model development, hyperparameter tuning and benchmarking; A.M. supervision and writing; M.H.S. supervision and writing. R.K.M., G.C. and H.C. have contributed equally. All authors have read and approved the manuscript.

