## Supplementary Figures and Tables for "miRXplain: explainable isomiR-aware microRNA target prediction using CLIP-L experiments and hybrid attention transformers"

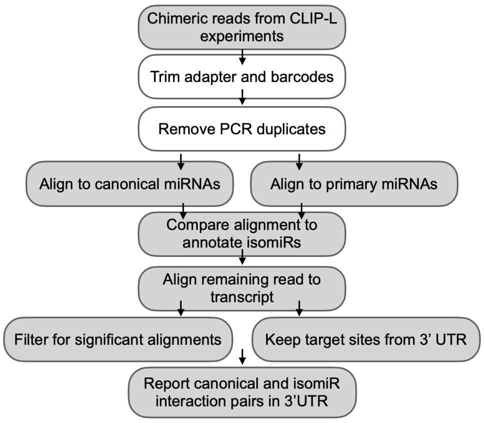

**Supplementary Figure 1**: Pipeline to retrieve canonical miRNAs and their variants ligated to their mRNA target sites from CLIP-L experimental chimeras. Gray-filled blocks denote processing steps added to the published SCRAP pipeline.

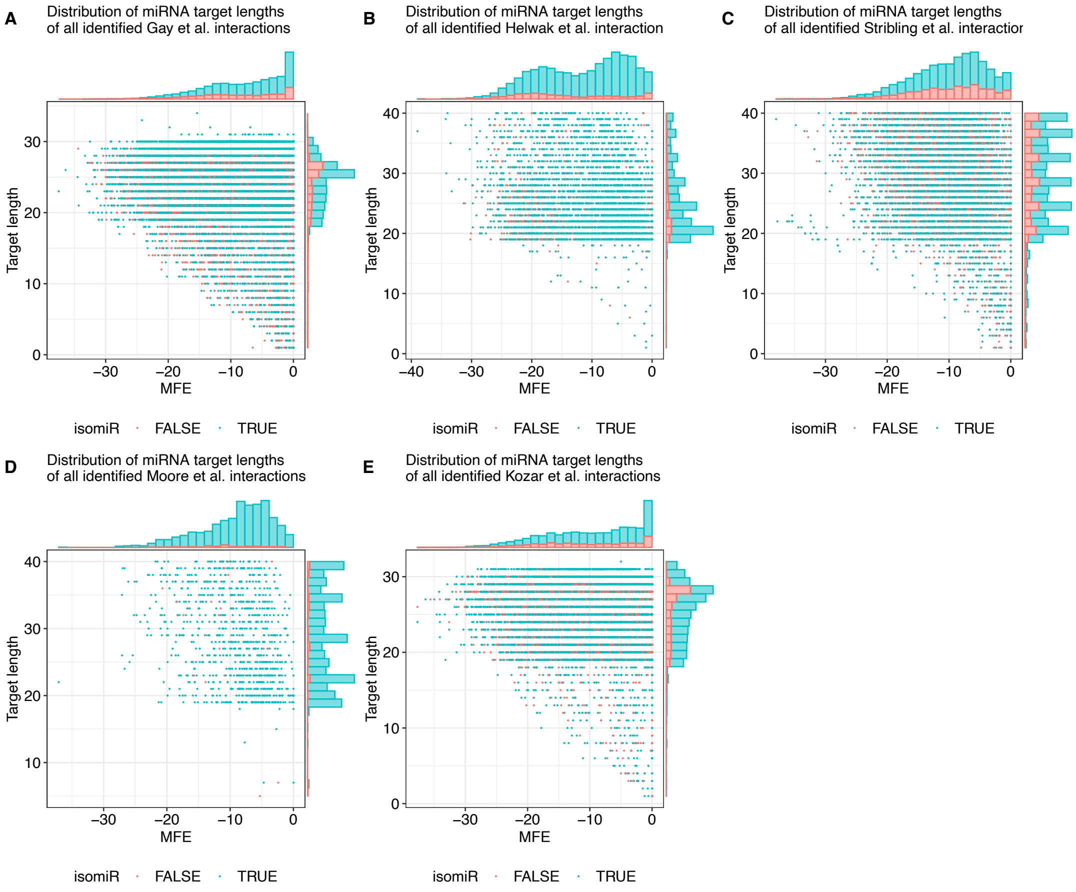

**Supplementary Figure 2**: Target length distributions over MFE for each CLIP-L experiment: (A) Gay et al. (B) Helwak et al. (C) Stribling et al. (D) Moore et al. and (E) Kozar et al. This analysis informed the choice of the minimum target-length threshold used prior to downstream analysis and modeling.

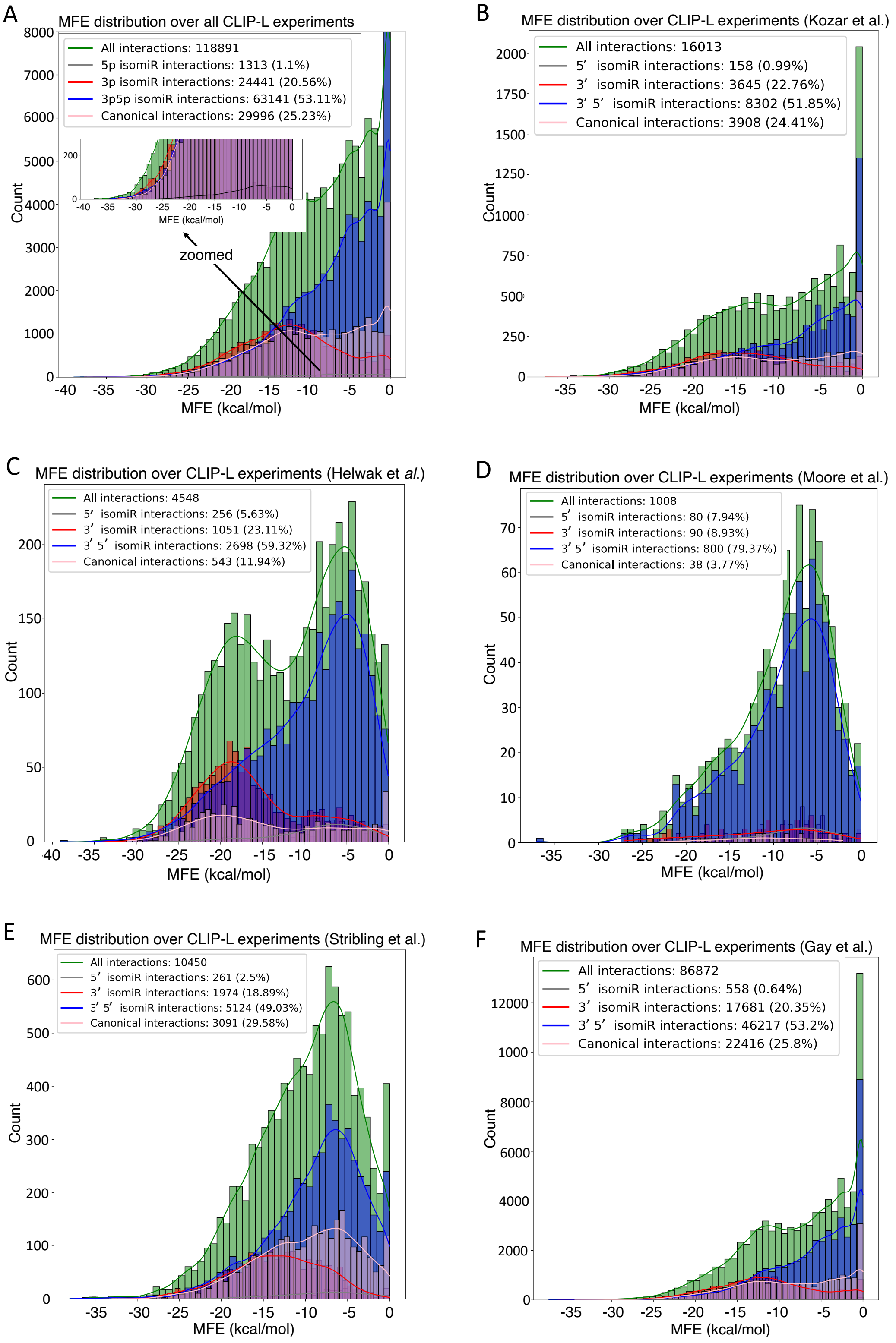

**Supplementary Figure 3**: Minimum free energy (MFE) distributions for all CLIP-L experiments indicate biases. MFE distributions for **(A)** all CLIP-L combined, and **(B)** Kozar et *al.* **(C)** Helwak et *al.* **(D)** Moore et *al.* **(E)** Stribling et *al.* and **(F)** Gay et *al.*

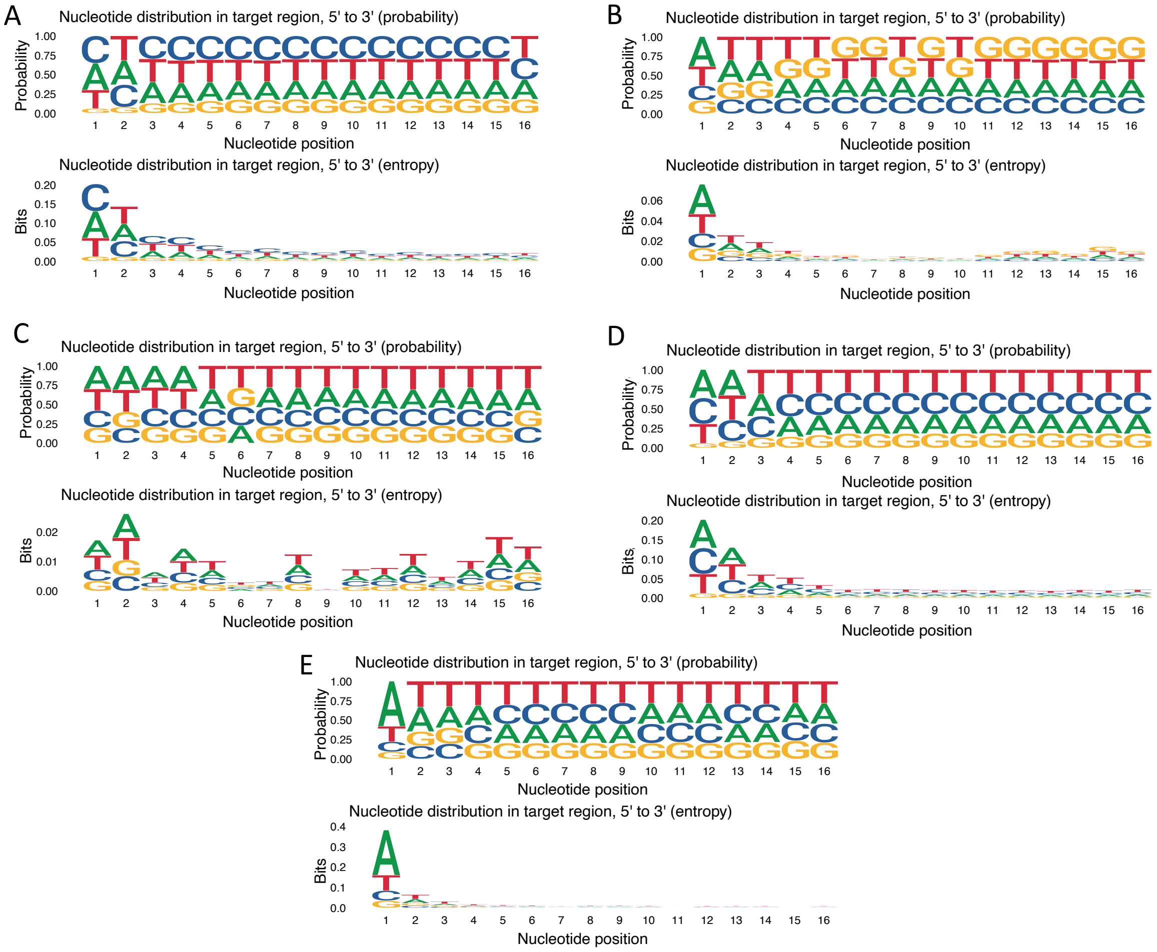

**Supplementary Figure 4**: Sequence logos of CLIP-L target sites showing nucleotide distributions for each experiment: **(A)** Kozar et al. **(B)** Helwak et al. **(C)** Moore et al. **(D)** Gay et al., and **(E)** Stribling et al.

**Supplementary Table 1**: Hyperparameter tuning search space used in *miRXplain*.

| **Hyperparameter** | **Range of Values** |
|---|---|
| Batch size | $\{32, 64, 128, 256, 512\}$ |
| Learning rate | $\{1 \times 10^{-3}, 1 \times 10^{-4}\}$ |
| Number of attention layers | $\{1, 2, 3\}$ |
| Number of attention heads | $\{1, 8\}$ |
| Model dimension | $\{64, 128, 256\}$ |
| FFN dimension | $\{64, 128, 256, 512\}$ |
| LBF rank | $\{32, 64, 128\}$ |
| Positional encoding dropout rate | $\{0, 0.1, 0.2\}$ |
| Attention dropout rate | $\{0.1, 0.2, 0.3\}$ |
| LBF dropout rate | $\{0, 0.1, 0.2, 0.3\}$ |
| CNN dropout rate | $\{0, 0.1, 0.2, 0.3\}$ |
| MLP dropout rate | $\{0, 0.1, 0.2\}$ |

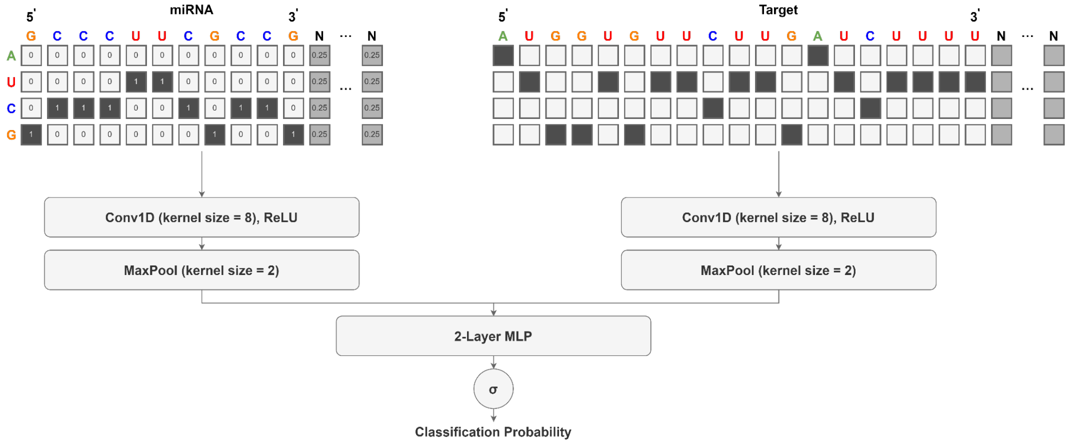

**Supplementary Figure 5**: CNN baseline architecture. The one-hot encoded miRNA and target sequences are separately processed through a convolutional layer followed by ReLU activation and max pool. Their output is concatenated and fed to a 2-layer MLP and a sigmoid function for the classification prediction.

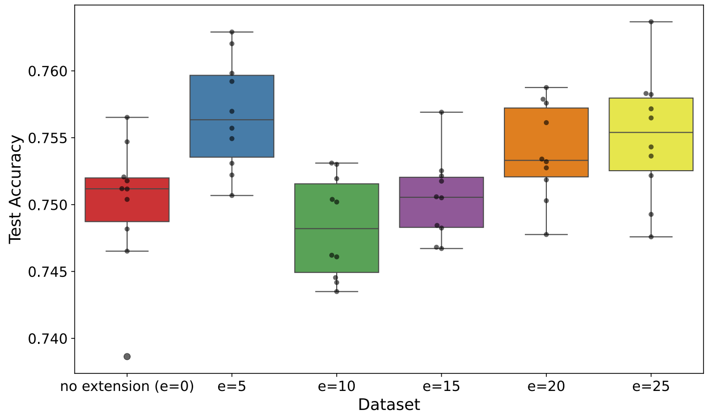

**Supplementary Figure 6**: Model classification accuracy using a *CNN baseline* on varying CLIP-L target lengths, extended towards the 3′ end of the target sites in intervals of 5 nucleotides.

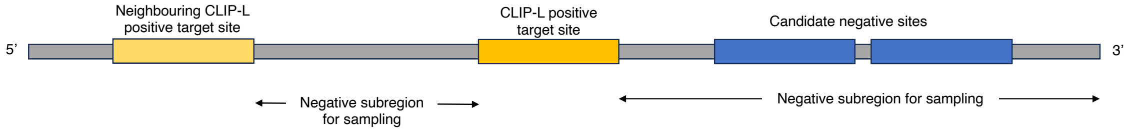

**Supplementary Figure 7**: Schema for negative sampling. The positive site refers to the CLIP-L reported target site. Candidate negatives were sampled from neighboring regions of the CLIP-L sites on the 3′ UTR, with MFE constraint and similar GC% as that of the positive.

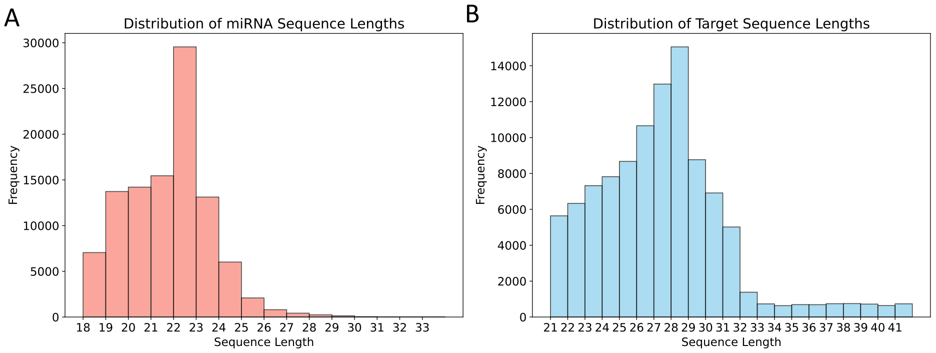

**Supplementary Figure 8**: Distribution of **(A)** miRNA and **(B)** target sequence lengths in our CLIP-L training dataset.

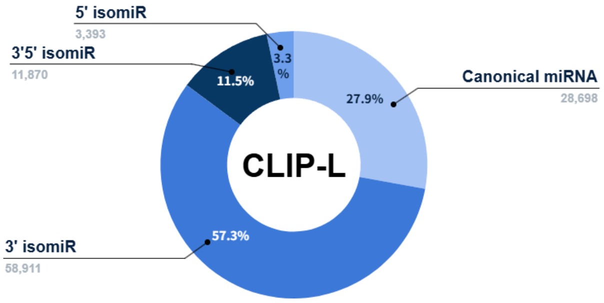

**Supplementary Figure 9**: Distribution of canonical and isomiR interactions in chimeras from CLIP-L experiments.

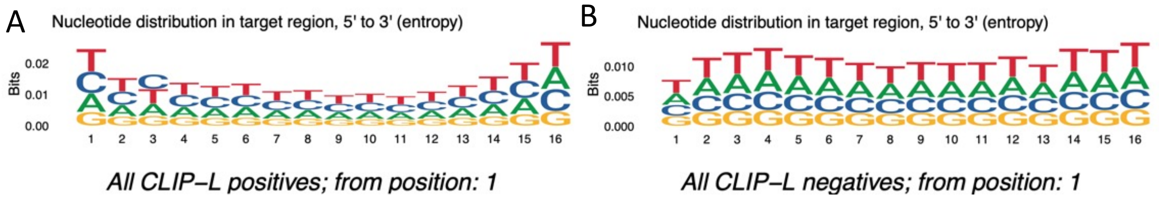

**Supplementary Figure 10**: Nucleotide distribution after bias correction, **(A)** over the positive sites from CLIP-L **(B)** over the negatives through sub sampling. The equal distribution ensure no bias on the model for any specific nucleotide at any position.

**Supplementary Table 2**: Top 5 miRXplain hyperparameter configurations

| # | Batch Size | LR | Attn Layers | Attn Heads | $d_{model}$ | $d_{ff}$ | LBF Rank | PE Dropout | Attn Dropout | CNN Dropout | MLP Dropout | Test Acc | Test Loss |
|---|---|---|---|---|---|---|---|---|---|---|---|---|---|
| 1 | 64 | 0.0001 | 2 | 1 | 64 | 512 | 32 | 0.0 | 0.3 | 0.3 | 0.0 | 0.8461 | 0.3539 |
| 2 | 32 | 0.0001 | 2 | 1 | 64 | 64 | 32 | 0.0 | 0.2 | 0.3 | 0.0 | 0.8463 | 0.3569 |
| 3 | 32 | 0.0001 | 3 | 1 | 64 | 256 | 32 | 0.2 | 0.3 | 0.2 | 0.1 | 0.8472 | 0.3571 |
| 4 | 128 | 0.0001 | 2 | 8 | 64 | 128 | 64 | 0.0 | 0.2 | 0.3 | 0.1 | 0.8411 | 0.3581 |
| 5 | 128 | 0.0001 | 1 | 8 | 128 | 512 | 64 | 0.2 | 0.2 | 0.1 | 0.1 | 0.8488 | 0.3596 |

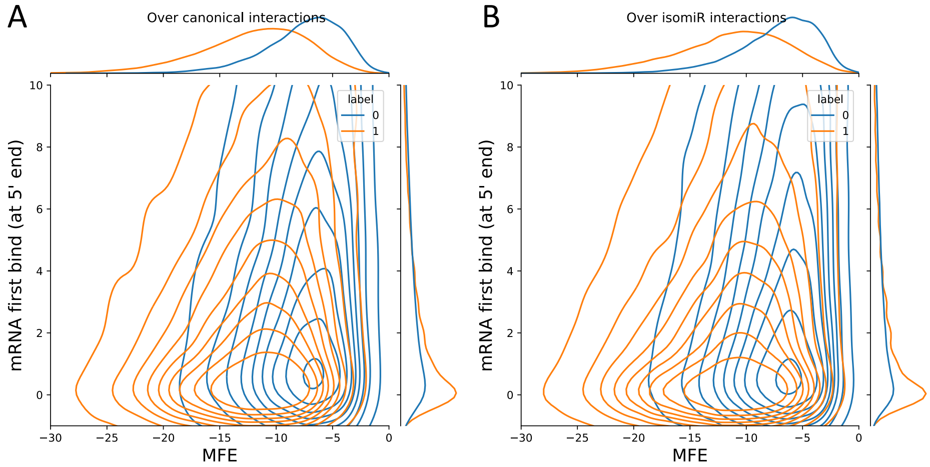

**Supplementary Figure 11**: Contour plots of MFE distributions in bias corrected positive miRNA target pairs (label 1) and among sampled negative interactions (label 0) for **(A)** canonical miRNA and **(B)** isomiR interactions. The positive interactions were from CLIP-L experiments, whereas the negatives were generated using the strategy described in Supplementary Fig. 7. The Y *axis* shows the first miRNA binding position along the mRNA (0-based) counting from the 5′ end. The X *axis* shows the MFE distributions in kcal/mol.

**Supplementary Table 3**: Formulas of extended classification metrics: accuracy, F1 score, TP: True Positives, FP: False positives, TN: True Negatives, FN: False Negatives.

| **Performance metric** | **Formula** |
|---|---|
| Accuracy | $\frac{TP+TN}{TP+TN+FP+FN}$ |
| F1 score | $\frac{2 \cdot \text{Precision} \cdot \text{Recall}}{\text{Precision} + \text{Recall}} = \frac{2TP}{2TP+FP+FN}$ |

**Supplementary Table 4**: Model parameters and training times in the ablation experiment of the bilinear fusion component.

| **Model** | **Model Parameters (M)** | **Training time (h)** |
|---|---|---|
| miRXplain (bilinear fusion) | **1.70** | **04:47** |
| miRXplain (concatenation) | 1.80 | 08:24 |
| miRXplain (subtraction) | 1.80 | 07:12 |

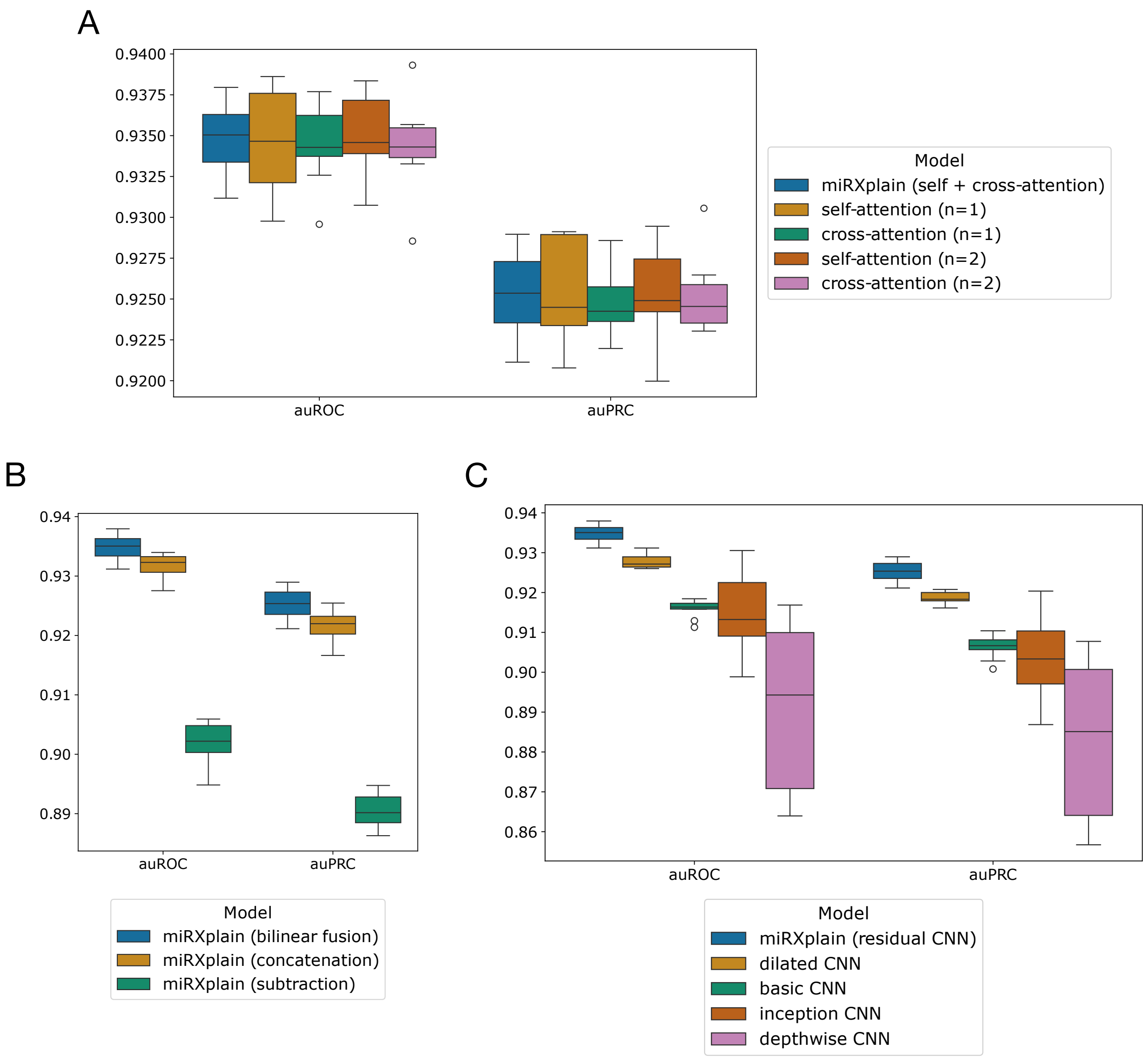

**Supplementary Figure 12**: Ablation study evaluating the attention type (**A**), the fusion module (**B**) and the type of refinement CNN employed (**C**). Panel **A** shows the developed model *miRXplain* with hybrid cross and self-attention, with only self-attention or only cross-attention, for a varying number of encoder layers (1 or 2) as specified by $n$ in brackets. Panel **B** shows *miRXplain* versus a model where the bilinear fusion component was either replaced by feature concatenation or element-wise subtraction of features. Panel **C** compares types of CNN architectures used for refinement, including *residual* (our model), *dilated*, *basic*, *inception* and *depthwise*.

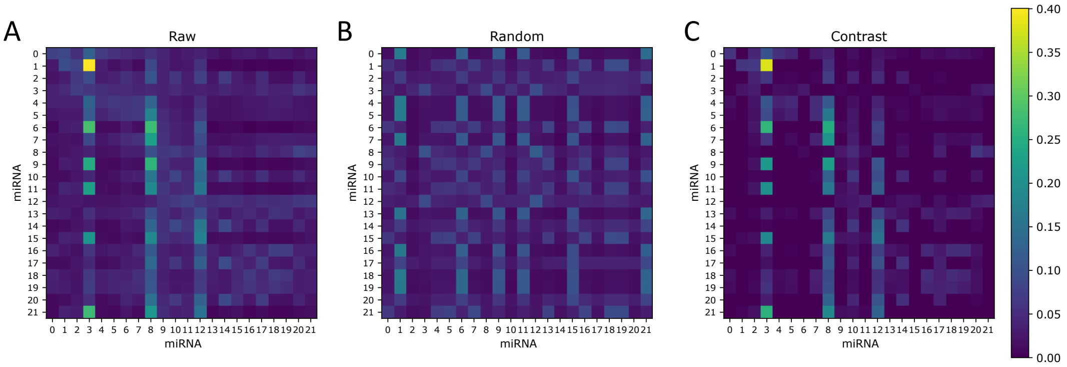

**Supplementary Figure 13**: Effects of correction of self-attention heatmaps with a background model, shown with miR-21-5p as example. **(A)** *Raw* is the original attention map, **(B)** *Random* is generated with a background model trained on a dataset with shuffled labels, and **(C)** *Contrast* is the background corrected map obtained by difference of *Raw* and *Random* maps, with clipping of negative values to 0.

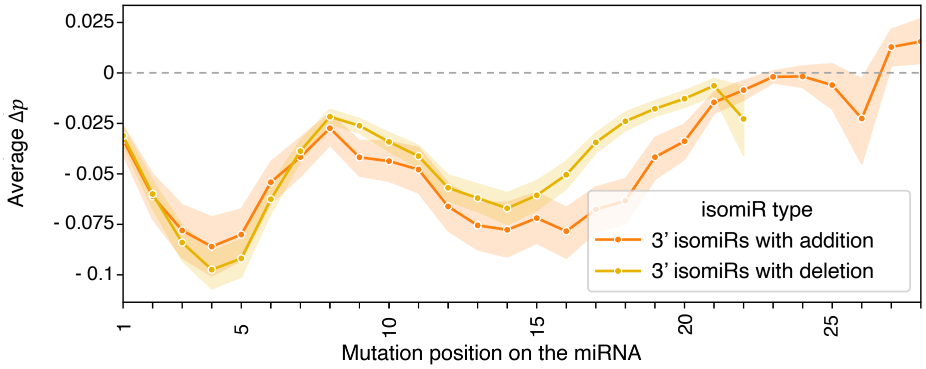

**Supplementary Figure 14**: Average prediction delta, $\Delta p$, $p_{\text{alt}} - p_{\text{ref}}$, as a function of the mutation position introduced along the 3′ isomiR sequences with addition and deletion events. Most 3′ isomiR addition and deletion events involve the addition or deletion of a single nucleotide at the 3′ end.

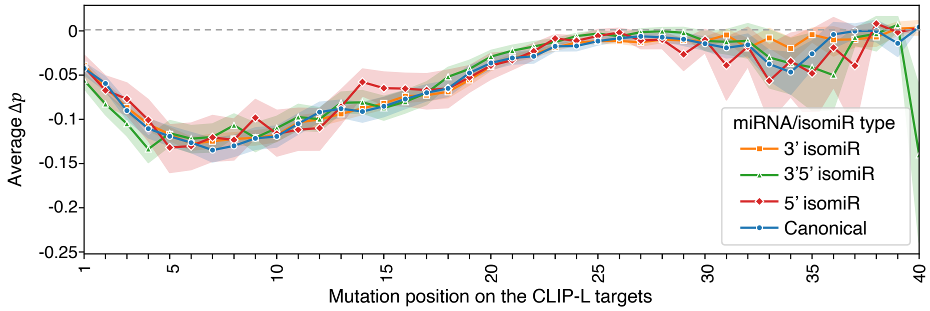

**Supplementary Figure 15**: Average prediction delta $\Delta p$, $p_{\text{alt}} - p_{\text{ref}}$ as a function of the mutation position introduced along the CLIP-L derived target sequences, for the 4 miRNA/isomiR types.

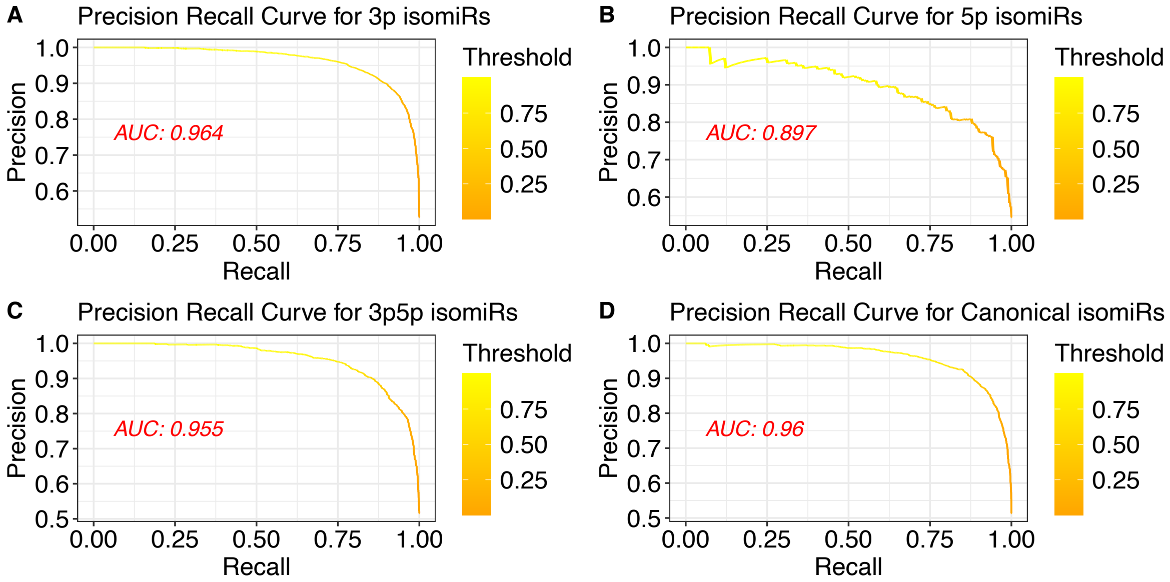

**Supplementary Figure 16**: PR curves *miRXplain* for **(A)** 3′ isomiRs **(B)** 5′ isomiRs **(C)** 3′5′ isomiRs, and **(D)** canonical miRNAs.

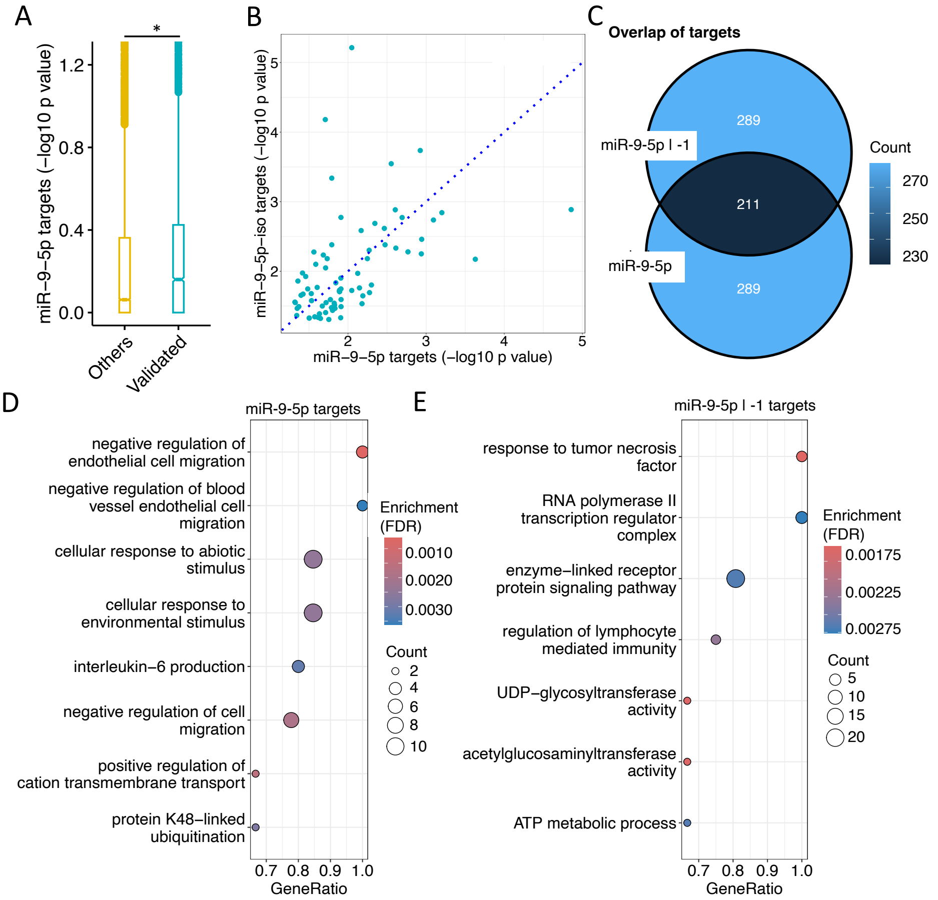

**Supplementary Figure 17**: *miRXplain* reveals miR-9-5p targets and associated functions. **(A)** Ranked p values obtained using miRXplain for miR-21-5p are significantly higher than those not validated. The significance test was performed using the Wilcoxon rank sum test, $p < 0.05$. *miRXplain* for **(B)** Comparison of gene $p$-values of canonical miR-9-5p (x-axis) against miR-9-5p isomiRs (y-axis), with -1 deletion events. Each dot denotes one target gene. **(C)** Overlap of canonical and isomiR targets of miR-9-5p. **(D,E)** Gene set functional enrichment results for **(D)** canonical targets, and **(E)**isomiRs.

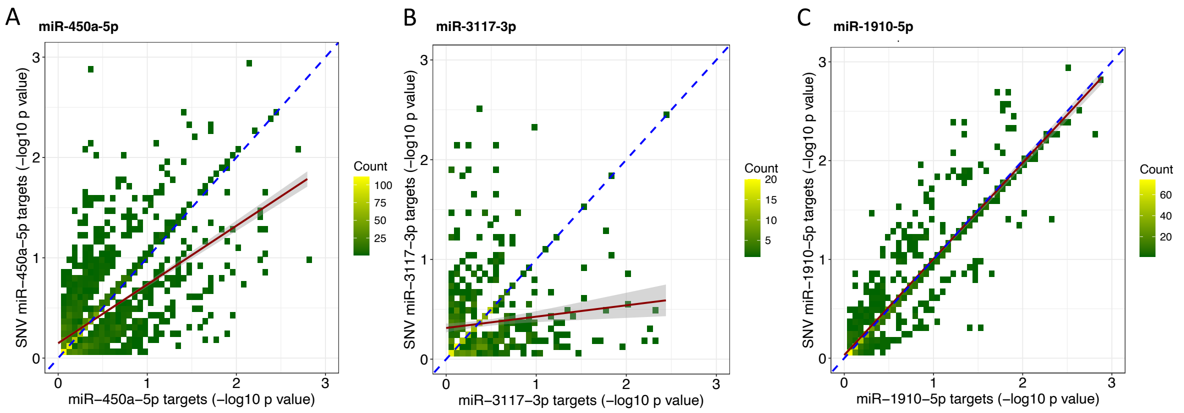

**Supplementary Figure 18**: *miRXplain* was used to investigate the effects of SNVs in miRNAs. Target gene binomial p values (workflow Fig. 6A) were computed for miRNAs overlapping SNVs. The binned scatter plots show the distribution of $p$ values for each validated target gene for (A) miR-450a-5p (B) miR-3117-3p, and (C) miR-1910-5p. The regression line (dark red), fitted using all target genes, indicates whether there is a consistent deviation from the diagonal (dashed blue).
